# Representations of evidence for a perceptual decision in the human brain

**DOI:** 10.1101/350876

**Authors:** Sebastian Bitzer, Hame Park, Burkhard Maess, Katharina von Kriegstein, Stefan J. Kiebel

## Abstract

In perceptual decision making the brain extracts and accumulates decision evidence from a stimulus over time and eventually makes a decision based on the accumulated evidence. Several characteristics of this process have been observed in human electrophysiological experiments, especially an average build-up of motor-related signals supposedly reflecting accumulated evidence, when averaged across trials. Another recently established approach to investigate the representation of decision evidence in brain signals is to correlate the within-trial fluctuations of decision evidence with the measured signals. We here report results for a two-alternative forced choice reaction time experiment in which we applied this approach to human magnetoencephalographic (MEG) recordings. These results consolidate a range of previous findings. In addition, they show: 1) that decision evidence is most strongly represented in the MEG signals in three consecutive phases, 2) that motor areas contribute longer to these representations than parietal areas and 3) that posterior cingulate cortex is involved most consistently, among all brain areas, in all three of the identified phases. As most previous work on perceptual decision making in the brain has focused on parietal and motor areas, our findings therefore suggest that the role of the posterior cingulate cortex in perceptual decision making may be currently underestimated.

## 1 Introduction

During perceptual decision making observers reason about the state of their environment. Supported by findings in single neurons of non-human primates, the underlying mechanism has been characterised as an accumulation-to-bound process (Gold & Shadlen, 2007). Specifically, the current consensus is that during perceptual decision making the brain accumulates noisy pieces of sensory evidence across time until it reaches a confidence bound. Most experimental results on this process have been based on stimuli which have been designed to provide the same amount of evidence per unit time on average across trials. Trial-averaged accumulated evidence then should follow a gradual build-up with evidence-dependent slope and a maximum close to the response within trial (Gold & Shadlen, 2007).

In humans, evidence of this kind of average build-up have been found using magneto- and electroencephalography (M/EEG). For example, lateralised oscillatory signals in the beta band measured with magnetoencephalography exhibit this build-up, where sources were located to dorsal premotor and primary motor cortex (Donner, Siegel, Fries, & Engel, 2009). In EEG, there are similar findings of a build-up for lateralised readiness potentials and oscillations (Kelly & O’Connell, 2013; de Lange, Rahnev, Donner, & Lau, 2013). Furthermore, when human participants have to detect the presence of stimuli in noise, a centro-parietal positivity shows the characteristics of an evidence-dependent build-up independently of the type of stimulus used and the kind of response made (Kelly & O’Connell, 2013; O’Connell, Dockree, & Kelly, 2012). Together these findings suggest that the human parietal and motor cortices are involved in perceptual decision making and in particular represent accumulated evidence. This view is compatible with electrophysiological recordings in non-human animals (Hanks & Summerfield, 2017) and an active role of the motor system during decision making (Cisek & Kalaska, 2010).

It has long been known that electromagnetic signals over motor areas build up towards a motor response and can signal an eventual choice even before the response (Smulders & Miller, 2012). This means that the crucial aspect of decision evidence representations is not the build-up as such, but its covariance with the theoretically available evidence. Consequently, more recent approaches have induced consistent, within-trial changes in available decision evidence (Wyart, de Gardelle, Scholl, & Summerfield, 2012; Thura & Cisek, 2014; Brunton, Botvinick, & Brody, 2013; Hanks & Summerfield, 2017). These within-trial changes allow more specific analyses, because one can directly assess the covariation between decision evidence and neural signals a) across a much richer sample of evidences than available with the trial-constant evidences in previous analyses and b) while the decision is ongoing.

Although it has previously been shown that electromagnetic signals in the human brain correlate with within-trial changing decision evidence (Wyart et al., 2012; de Lange, Jensen, & Dehaene, 2010; Gluth, Rieskamp, & Büchel, 2013; Gould, Nobre, Wyart, & Rushworth, 2012), these studies had either rather long stimulus presentation times atypical for fast perceptual decisions (Gluth et al., 2013; Gould et al., 2012), or did not employ a reaction time paradigm (Wyart et al., 2012; de Lange et al., 2010; Gould et al., 2012). In the present work we therefore sought representations of decision evidence in a two-alternative forced choice reaction time paradigm in which we induced changes in decision evidence every 100 ms. That is, our paradigm attempts to mimic natural perceptual decision making behaviour more closely than previous investigations with controlled, within-trial changing evidence while still observing neural responses across the whole human brain.

Specifically, we investigated correlations between decision evidence and human MEG signals and their sources. We found particularly large effects of decision evidence in the human MEG in three consecutive phases aligned to when the particular piece of evidence became available. The underlying sources indicate that the information delivered by the evidence propagated from visual over parietal to motor areas, as expected, but also that it remained in motor areas for a longer time than in parietal areas. In addition, our results implicate posterior cingulate cortex in all of the identified phases suggesting a central role of this brain region in the transformation of sensory signals to decision evidence in our task.

## 2 Results

While MEG was recorded, 34 human participants observed a single white dot on the screen changing its position every 100 ms and had to decide whether a left or a right target (two yellow dots) was the centre of the white dot movement (Figure 1). Under moderate time pressure (see Methods), participants indicated their choice with a button press using the index finger of the corresponding hand. The distance of the target dots on the screen was chosen in behavioural pilots so that participants had an intermediate accuracy around 75% while being told to be as accurate and fast as possible. The average median response time across participants was 1.1 s with an average accuracy of 78% (cf. Figure 1B).

**Figure 1:**
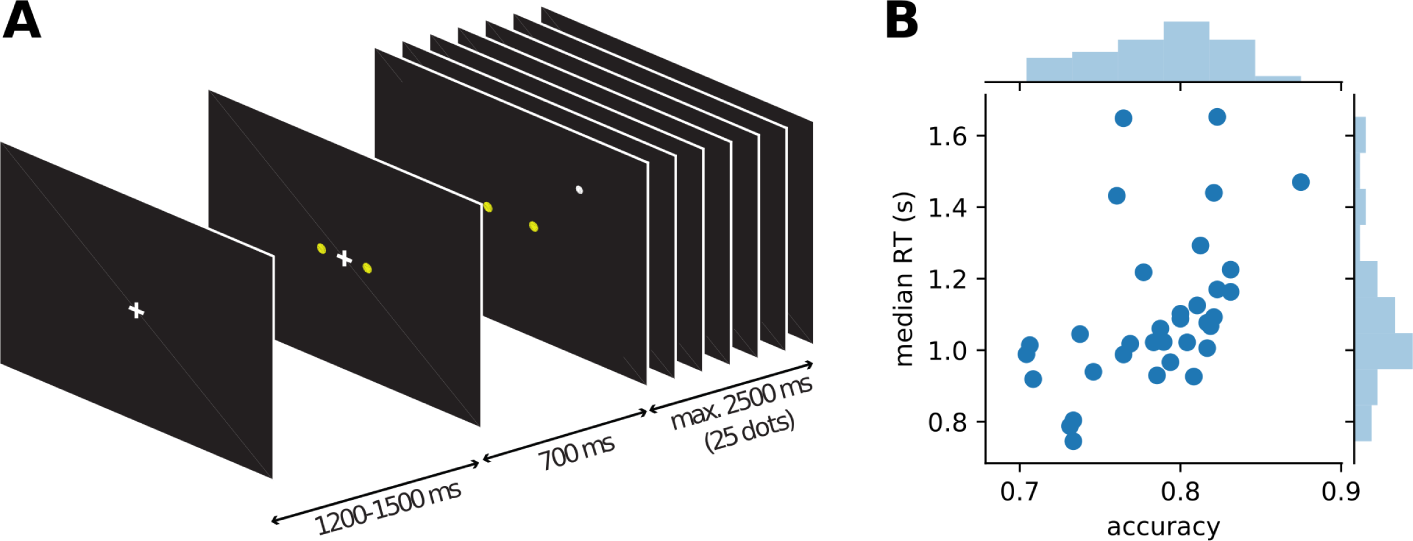
Course of events within a trial in the single dot task (A) and behaviour of individual participants (B). Each trial started with the presentation of a fixation cross, followed by the appearance of the two yellow targets after about 1 s. 700 ms after the appearance of the targets the fixation cross disappeared and a single white dot was presented at a random position on the screen (drawn from a 2D-Gaussian distribution centred on one of the targets). Every 100 ms the position of the white dot was changed to a new random draw from the same distribution. Participants were instructed to indicate the target which they thought was the centre of the observed dot positions. After 25 dot positions (2.5 s) without a response, a new trial was started automatically, otherwise a new trial started with the response of the participant. Average behaviour (accuracy and median response time) for each of the 34 participants is shown in B.

This paradigm dissociates two different kinds of information available to the participants from the stimulus. The x-coordinates of the jumping white dot convey decision-relevant perceptual information while the y-coordinates convey perceptual information that is irrelevant for the decision. We assume that both signals are processed by the brain, but only the decision-relevant x-coordinates are taken into account when making a decision.

To define decision evidence, we used a computational model. An ideal observer model for inference about the target given a sequence of single dots has been described before (H. Park, Lueckmann, von Kriegstein, Bitzer, & Kiebel, 2016; Bitzer, Park, Blankenburg, & Kiebel, 2014). This model identifies, as expected, the x-coordinates of the white dot positions as momentary decision evidence. Specifically, there is a direct linear relationship between x-coordinates and momentary evidence so that in the following regression analyses we could directly use the x-coordinates as independent variables instead of having to compute decision evidence from the x-coordinates through the model. We further identified the cumulative sum of x-coordinates across single dot positions as accumulated evidence which corresponds to the average state of a discrete-time drift-diffusion model (Bitzer et al., 2014).

### 2.1 Participants integrate evidence provided by single dot positions to make decisions

As the task required and the model predicted, participants made their decision based on the provided evidence. In Figure 2 we show this as the correlation of participants’ choices with momentary and accumulated evidence. Momentary evidence was mildly correlated with choices throughout the trial (correlation coefficients around 0.3) while the correlation between accumulated evidence and choices increased to a high level (around 0.7) as more and more dot positions were presented. This result indicates that participants accumulated the momentary evidence, here the x-coordinate of the dot, to make their choices. In contrast, as expected, the y-coordinates had no influence on the participants’ choices as indicated by correlation coefficients around 0 (Figure 2B).

**Figure 2:**
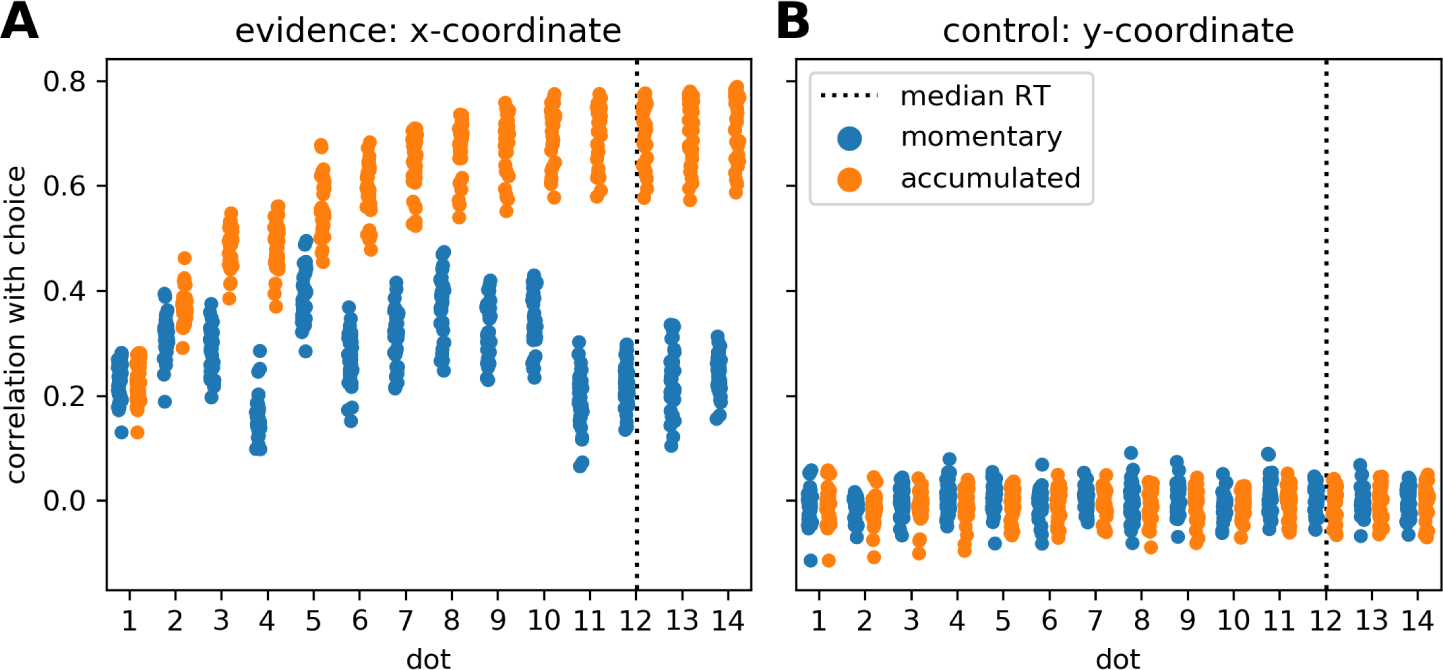
Participants accumulate momentary evidence provided by dot positions for making their decisions. (A) Each shown point corresponds to the Pearson correlation coefficient for the correlation between choices of a single participant and the sequence of presented dot positions across the 480 trials of the experiment. We plot, over stimulus duration, the momentary (blue) and the accumulated evidence (orange). The dotted vertical line shows the median RT across participants. Until about the 10th dot presentation the correlation between accumulated evidence and participant choices rises, reaching values around 0.7 while the momentary evidence is only modestly related to participant choices across all dots. (B) The same format as in A but all measures are computed from the y-coordinates of dot positions which were irrelevant for the decision. As expected, y-coordinates do not correlate with participant choices.

Previous work has investigated the influence of individual stimulus elements on the eventual decision and whether this influence differed across elements (Wyart et al., 2012; Hubert-Wallander & Boynton, 2015). In our analysis this corresponds to checking whether the correlations with momentary evidence shown in Figure 2A differ across dots. This is clearly the case (*F*(13,462) = 65.49,*p* ≪ 0.001). Contrary to previous work (Hubert-Wallander & Boynton, 2015) we do not observe a primacy effect. Instead, we observe a particularly large difference in the influence of the 4th and 5th dots on the decision (post-hoc paired t-test: *t*(33) = —34.90, *p* ≪ 0.001) with the 5th dot having a strong influence while the 4th dot having a relatively small influence. This reflects our pre-selection and manipulation of stimuli which were partially chosen from a previous experiment to induce large response times (leading to the small influence of the 4th dot) and a manipulation of the 5th dot to create large variation in x-coordinates (see Methods for further details). Taken together these results confirm that the used stimuli were effective in driving the decisions of the participants and that the theoretically defined momentary and accumulated evidence integrate well with observed behaviour.

### 2.2 MEG signals covary with momentary evidence at specific time points after stimulus update

For the analysis of the MEG data we used regression analyses computing event-related regression coefficients of a general linear model (Clarke, Taylor, Dev-ereux, Randall, & Tyler, 2013; Hauk, Davis, Ford, Pulvermuller, & Marslen-Wilson, 2006). For our main analysis the regressors of interest were the momentary evidence and, as a control, the y-coordinates of the presented dots. We normalised both the regressors and the data so that the resulting regression coefficients can be interpreted as approximate correlation values while accounting for potential covariates of no interest (see Methods). Note that this correlation analysis contrasts with standard event-related field analyses, where one would only test for the presence of a constant time-course across trials. With the correlation analysis, the estimated regression coefficients describe how strongly the MEG signal, in each time point and each sensor (or source), followed the ups and downs of variables such as the momentary evidence, across trials.

As a first result, we found that correlations between momentary evidence and MEG signals followed a stereotypical temporal profile after each dot position update (cf. Supplementary Figure 1). Therefore, we performed an expanded regression analysis where we explicitly modelled the time from each dot position update, which we call ‘dot onset’ in the following. To exclude the possibility that effects signalling the button press motor response influence the results of the dot onset aligned analysis, we only included data, for each trial, up until at most 200 ms prior to the participant’s response.

We first identified time points at which the MEG signal correlated most strongly with the momentary evidence. For these sensor-level analyses we focused on magnetometer sensors only. We performed separate regression analyses for each time point from dot onset, magnetometer sensor and participant, computed the mean regression coefficients across participants, took their absolute value to yield a magnitude and averaged them across sensors. Figure 3 shows that the strongest correlations between momentary evidence and magnetometer signals occurred at 120 ms, 180 ms and in a prolonged period from roughly 300 to 500 ms after dot onset. In contrast, correlations with the decision irrelevant control variable, that is, the dot y-coordinates, were significantly lower in this period from 300 to 500 ms (two-tailed Wilcoxon test for absolute average coefficients across all sensors and times within 300-500 ms, *W* = 382781,*p* ≪ 0.001).

**Figure 3:**
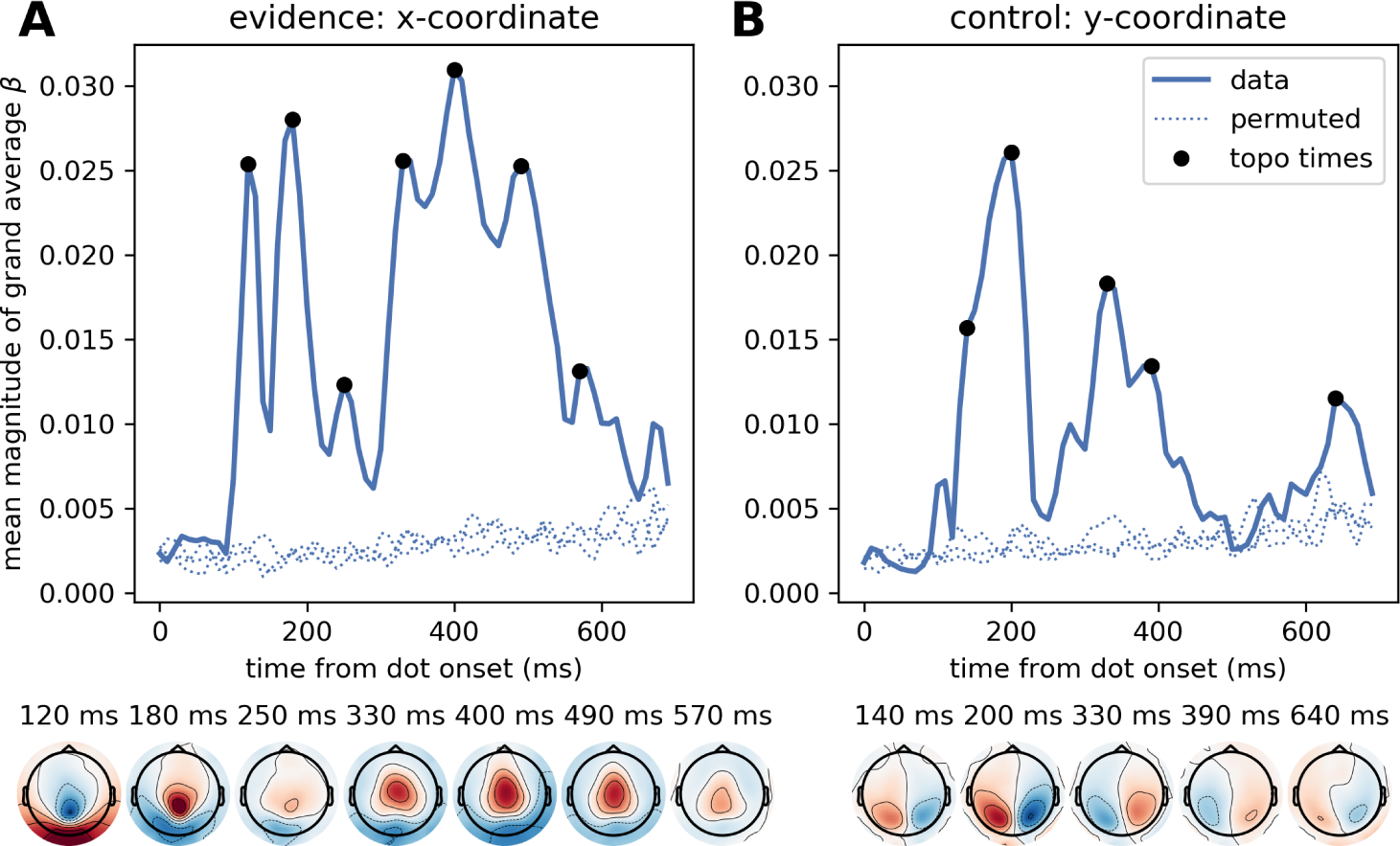
Time course of correlation strengths between magnetometer measurements and momentary evidence (left) and perceptual control variable (right). Top panels show time courses of the mean (across sensors) magnitude of grand average regression coefficients (*β*). For comparison, dotted lines show the corresponding values for data which were randomly permuted across trials before statistical analysis. Black dots indicate time points for which the sensor topography is shown below the plot. These topographies directly display the grand average regression coefficients at the indicated time without rectification, i.e., with negative (blue) and positive (red) correlation values. (A) The momentary evidence has strong correlations with the magnetometer signal at 120 ms, 180 ms and from about 300 ms to 500 ms after dot onsets. (B) The correlations with the decision irrelevant y-coordinate are visibly and significantly weaker than for the evidence, but there are two prominent peaks from about 120 ms to 210 ms and at 320 ms after dot onset. There is no sustained correlation with the y-coordinate beyond 400 ms and the topographies of magnetometers differ strongly between evidence and y-coordinates. Specifically, the evidence exhibits occipital, centro-parietal and central topographies whereas the y-coordinate exhibits strong correlations only in lateral occipito-parietal sensors.

The sensor topographies shown in Figure 3 indicate for the momentary evidence a progression of the strongest correlations from an occipital positivity over a centro-parietal positivity to a central positivity. y-coordinate correlations, on the other hand, remained spatially at occipito-parietal sensors.

### 2.3 Correlations with accumulated evidence

Guided by the model we used dot x-coordinates as representation of momentary evidence, but dot x-coordinates also do have a purely perceptual interpretation similar to the y-coordinates as they simply measure the horizontal location of a visual stimulus. Correlations with x-coordinates, therefore, may reflect at some time points early visual processes independent of the decision, at some time points momentary evidence and other time points both of them. Contrasting the strength of significant effects for x- and y-coordinates (Figure 3) already suggested that at least from 400 ms after dot onset x-coordinates indeed represented momentary evidence. To further corroborate this supposition we turned to a form of decision evidence that has no direct purely perceptual interpretation and is more closely related to the decision itself: the accumulated evidence.

Note that accumulated evidence is, through the final choice, more strongly related to the motor response than the momentary evidence (cf. Figure 2A Supplementary Figure 2 which means that some effects indicated by the accumulated evidence regressor may be attributed to the motor response and not the accumulated evidence. To account for this potential confound we excluded also from this analysis all data later than 200 ms before the response so that the results only contain effects unrelated to the motor response.

Furthermore, accumulated and momentary evidence are themselves entangled such that both regressors lead to partially overlapping effects. See Methods and Supplementary Material for more information. The point of this analysis, however, is that it will more strongly highlight accumulated evidence effects while the momentary evidence regressor in the previous analysis more strongly highlighted perceptual and momentary evidence effects.

Figure 4 depicts the time course of overall correlation magnitudes for accumulated evidence together with effect topographies at chosen time points. We found correlations between the MEG signal and accumulated evidence at all of peri-stimulus time until about 550 ms after dot onset. Crucially, at all time points of that period we observed centro-parietal and, especially, central sensor topographies suggesting that these represent specifically decision-relevant information such as momentary or accumulated evidence, as hypothesised based on the correlations with x-coordinates shown in Figure 3. For further discussion of the time course of accumulated evidence, see Supplementary material.

**Figure 4:**
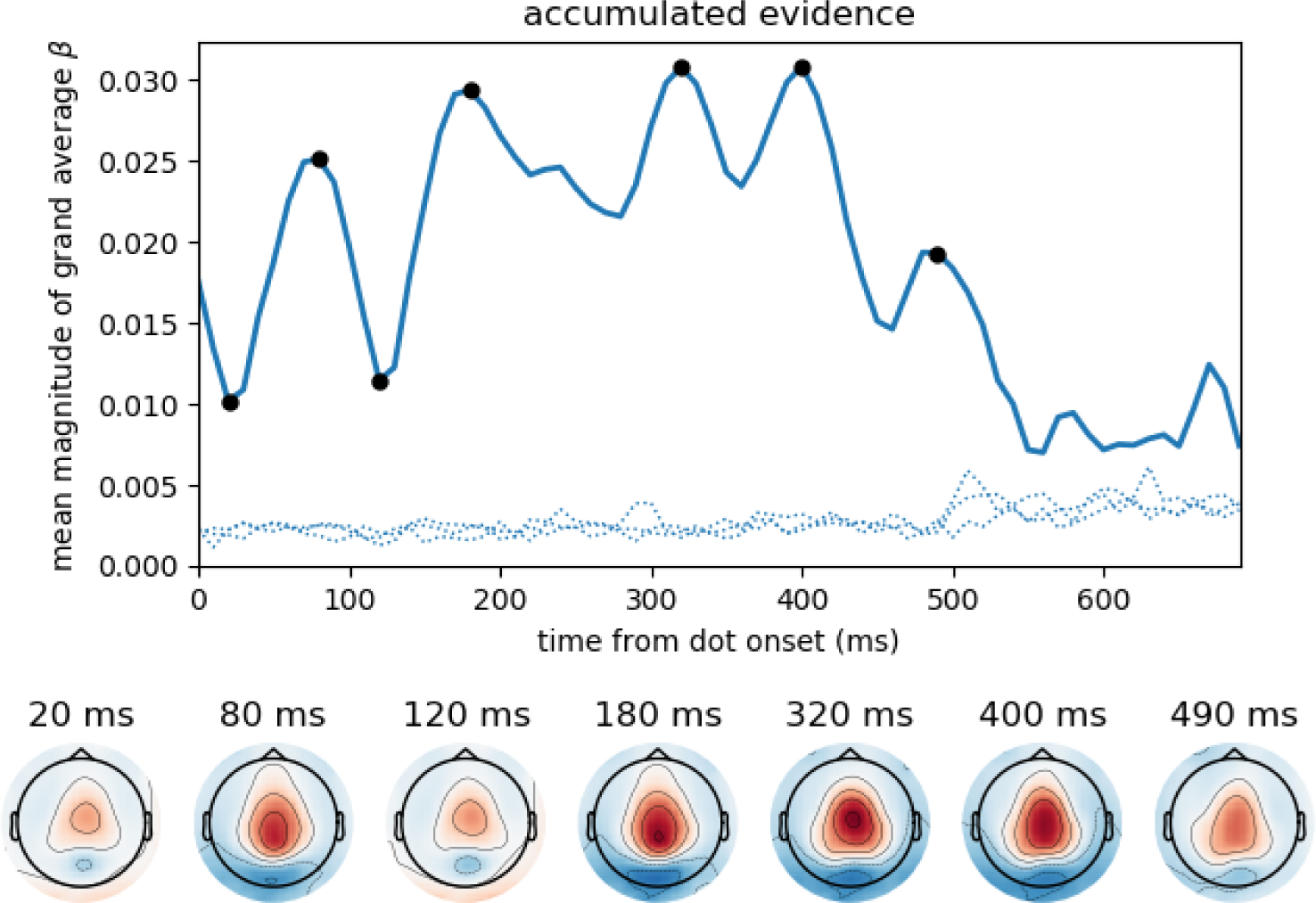
Accumulated evidence correlated with magnetometer signals from 0 to about 550 ms after dot onset displaying central sensor topographies throughout this time period. Format as in Figure 3, i.e., top panel shows time course of the mean (across sensors) magnitude of grand average regression coefficients (*β*) together with corresponding time courses after 3 different permutations across trials (dotted). For further analysis and discussion of results, see Supplementary Material.

### 2.4 Sources of stimulus-aligned momentary evidence effects

By investigating the sources of the evidence correlations at sensor level, we aimed to better understand the nature of these effects and to confirm their locations in the brain suggested by the shown sensor topographies. In particular, we were interested in linking the time points at which we found strong momentary evidence correlations to potential functional stages in the processing of decision evidence, such as sensory processing, relating sensory information to the decision and integrating momentary evidence with previous evidence.

We reconstructed source currents along the cerebral cortex for each participant and subsequently repeated our regression analysis on the estimated sources. Specifically, we performed source reconstruction on the preprocessed MEG data using noise-normalised minimum norm estimation based on all MEG sensors (magnetometers and gradiometers) (Gramfort et al., 2014, 2013; Dale et al., 2000). Further, we aggregated estimated values by averaging across sources within 180 brain areas defined by a recently published brain atlas (Glasser et al., 2016). This resulted in average time courses for each experimental trial in each of the 180 brain areas defined per hemisphere for each participant. We then applied the expanded regression analysis to these source-reconstructed time courses instead of onto MEG sensors. Following the summary statistics approach we identified time points and areas with significant second-level correlations by performing t-tests across participants and applying multiple comparison correction using false discovery rate (Benjamini & Hochberg, 1995) simultaneously across all time points and brain areas.

The time course of correlation magnitudes shown in Figure 3 suggested three time windows at which particularly strong correlations with momentary evidence were present in the brain. The source analysis gives equivalent results: Multiple comparison corrected effects occurred only within 110 ms – 130 ms, 160 ms – 200 ms and 290 ms – 510 ms (cf. Source Data 1). In subsequent analyses we, therefore, concentrated on these time windows and call them according to their temporal order “early”, ‘’intermediate” and “late” phases. Figure 5 depicts the brain areas with at least one significant multiple comparison corrected effect within the corresponding phase. The colour scale indicates the average t-value magnitudes within the time window for these significant areas (we chose to display t-value magnitudes instead of correlation magnitudes here, because the estimated correlation values had larger second-level variability differences across brain areas than sensors).

**Figure 5:**
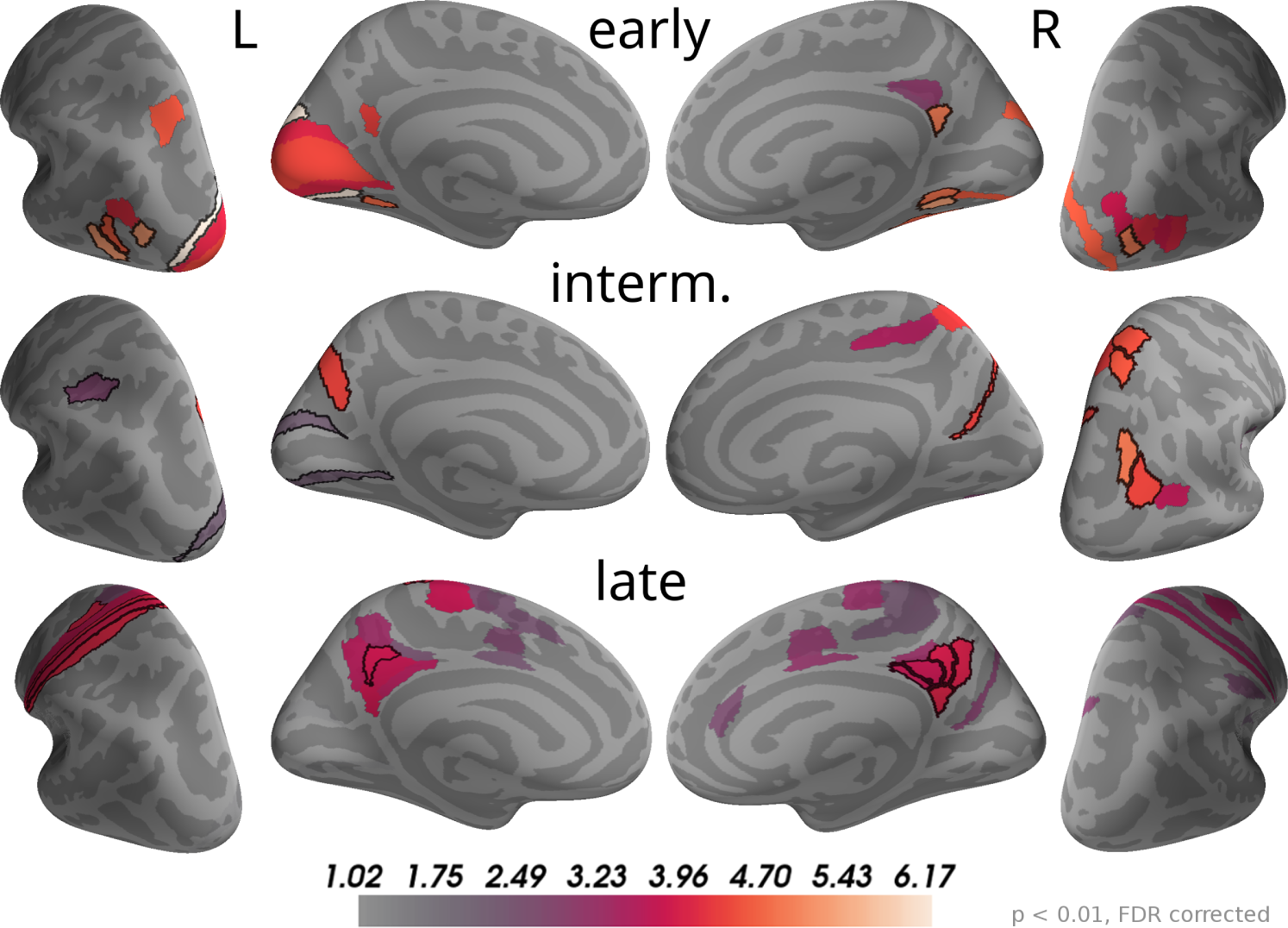
Correlations with momentary evidence shift from visual over parietal to motor and posterior cingulate areas. We investigated the three time windows with strong correlations in the sensor-level results: early (110 ms – 130 ms), intermediate (160 ms – 200 ms) and late (290 ms – 510 ms). For each of these phases only brain areas with at least one significant effect (*p* < 0.01, FDR corrected) within the time window are coloured. For display purposes, colours show average second-level t-value magnitudes where the average is taken over time points within the time window. The 5 areas with the most consistent, strong correlations per hemisphere and time window are marked by black outlines. These were (in that order; specified as Brodmann areas with subdivisions as defined in (Glasser et al., 2016)): early, left – V3, FST, LO3, VMV2, MST; right – VMV2, LO1, v23ab, VMV1, VVC; intermediate, left – POS2, AIP, V2; right – IP0, VIP, 7AL, PGp, DVT; late, left – 1, 3a, 6d, 3b, 31pd; right – v23ab, 7m, 31pd, 31pv, d23ab.

As the sensor topographies suggested, we observed that in the early phase the strongest correlations were located in visual areas such as V3, V1 and areas in the lateral occipital cortex (e.g., FST, MST, LO3 according to (Glasser et al.,2016)), but also in a small area of posterior cingulate cortex (v23ab) and there was an effect in a parietal area of the left hemisphere (MIP). In the intermediate phase most of the correlations in visual areas, especially those in lateral occipital areas, vanished. Instead, more parietal areas exhibited significant correlations with momentary evidence, especially in the right inferior (IP0, PGp) and superior parietal cortex (VIP, 7AL, 7Am). Additionally, we found strong correlations in posterior cingulate cortex (POS2 and DVT). In the late phase some correlations in parietal areas persisted, but only focal at some time points so that on average across the time window correlations were weak compared to other brain areas. Specifically, the strongest correlations were spread across the posterior cingulate cortex in both hemispheres (especially areas v23ab, 31pd, 7m, 31pv, d23ab). Further strong correlations occurred in motor areas, especially in the left hemisphere, including somatosensory areas (3a, 3b, 1), primary motor cortex (area 4) and premotor areas (6a, 6d). Note that we excluded from the analysis all time points later than 200 ms before the trial-specific motor response. Additionally, we observed weaker correlations in mid and anterior cingulate motor areas (e.g., 24dv, p24pr). These results confirm that the information carried by the decision-relevant x-coordinates shifts from visual over parietal areas towards motor areas where this information, presumably momentary evidence, appears to be represented over a longer time period. The results also reveal that source currents of brain areas in posterior cingulate cortex had strong correlations with x-coordinates throughout all three phases. Accordingly, the areas with the largest correlation magnitudes on average across all time points within 0 to 500 ms were predominantly located in posterior cingulate cortex (5 areas with strongest average effects in that order: left – v23ab, 3a, 31pd, 3b, 1; right – v23ab, DVT, d23ab, 31pv, 7m). This suggests a potentially central role of posterior cingulate cortex in the processing of momentary evidence in the task.

### 2.5 Sources of stimulus-aligned accumulated evidence effects

The sensor topographies for the accumulated evidence effects suggested that accumulated evidence was represented in common brain sources across the whole time window of 0 to 550 ms from dot onset. Therefore, we used this full time window to investigate the underlying sources. As for the momentary evidence, cf. Figure 5 we identified brain areas with significant correlations after FDR correction across locations and times (*p* < 0.05, no significant effects for *p* < 0. 01) in at least one time point and then averaged the t-value magnitudes across time points within the time window in these areas. Given the similarity of sensor topographies of momentary evidence in the late phase and the sensor topographies of accumulated evidence we expected their sources to overlap.

In Figure 6 one can see that, although the estimated correlation magnitudes were slightly higher for the accumulated evidence than for the momentary evidence, fewer effects were statistically significant for accumulated evidence. This is most likely because the variability of correlation magnitudes across participants increased relative to momentary evidence effects (results not shown). Otherwise, the identified brain areas were consistent with those of the momentary evidence in the late phase. In particular, we observed consistently strong correlations with accumulated evidence in motor, premotor, cingulate motor and posterior cingulate areas.

**Figure 6:**
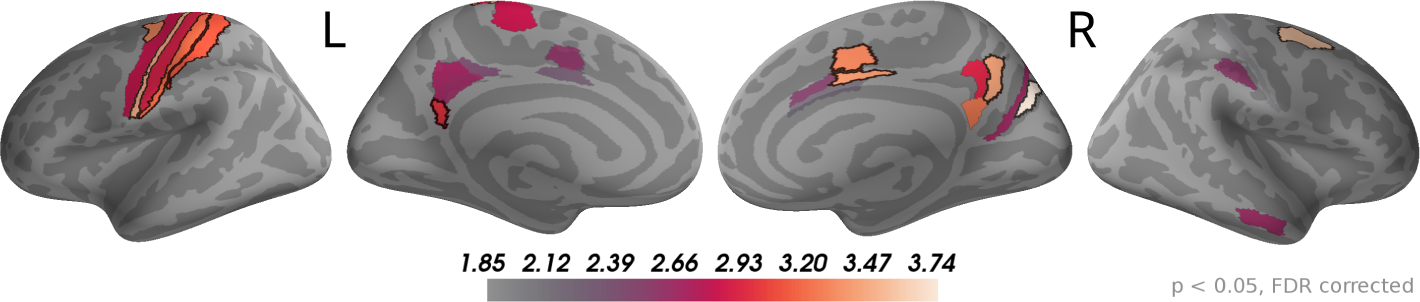
Sustained correlations with accumulated evidence in motor and cingulate areas. Following the procedure in Figure 5, we coloured only areas with a significant correlation with accumulated evidence (*p* < 0.05 FDR corrected) with colour indicating the average t-value magnitude in the extended time window from 0 ms to 550 ms after dot onset. The 5 largest effects were (marked by black boundaries): left – 3a, 6d, 1, 2, v23ab; right – V6, 6a, 7m, p24pr, 24dv.

### 2.6 Correlations with choice reveal response-aligned buildup and separate motor response

Our finding that momentary or accumulated evidence is represented in motor areas is consistent with a wide range of previous work (Donner et al., 2009; Kelly & O’Connell, 2013; de Lange et al., 2013; Thura & Cisek, 2014; Selen, Shadlen, & Wolpert, 2012; Michelet, Duncan, & Cisek, 2010). If motor areas are involved in processing momentary or accumulated evidence prior to a response, as these results indicate, the question arises how these processes relate to motor processes linked to the response itself. More specifically, we were interested in how the patterns of correlations with momentary and accumulated evidence related to correlation patterns representing the motor response and whether these could be linked to the absence or presence of the involvement of certain brain areas. To investigate correlation patterns representing the motor response we computed choice-dependent effects centred on the response time of the participants. We did this with a regression analysis using the participant choice as a regressor of interest (see Methods). The choice regressor provides a measure for how well the choice of the participants can be decoded from univariate brain signals.

Figure 7 depicts the estimated time course of correlation magnitudes averaged across participants and sensors. From about 500 ms before the response, correlations between choice and MEG data became gradually stronger culminating in an expected peak centred slightly after the response. The sensor topographies of the build-up period before the response strongly resembled those we found for accumulated evidence in our previous analyses. In fact, these results most likely correspond to the same effect, because the participant choice itself was increasingly correlated with accumulated evidence as the trial progressed (cf. Figure 2). That is, the build-up seen in the figure only indirectly visualises an increasing evidence signal by depicting an increasing alignment of the final choice with the internal representation before the response (presumably accumulated evidence).

**Figure 7:**
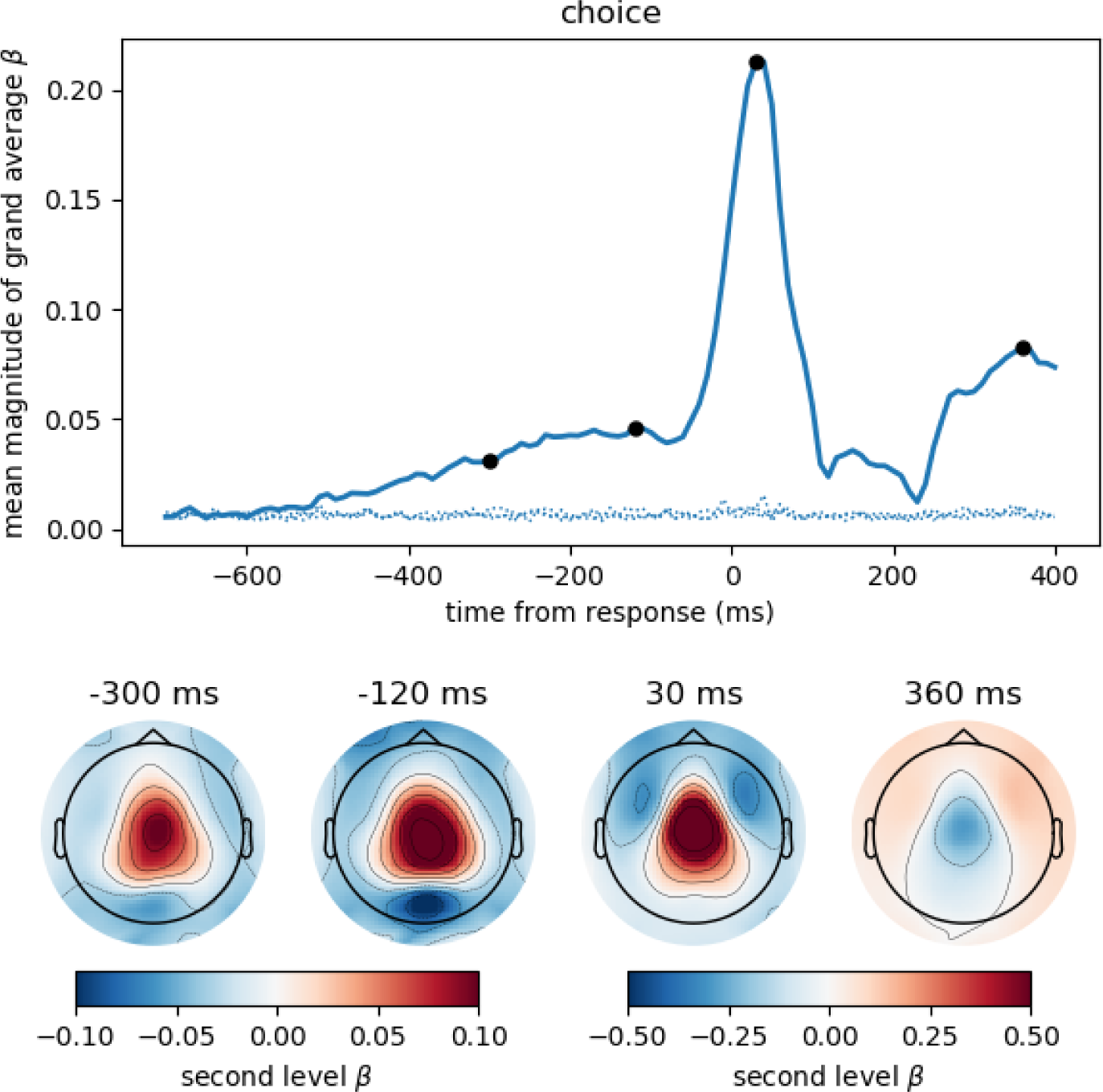
The button press motor response is also represented most strongly in central magnetometers, but the corresponding topography differs slightly from that associated with momentary and accumulated evidence. We computed the correlation between participant choices and MEG magnetometers using linear regression for data aligned at response time. Following the format of Figure 3 we here show the time course of the mean (across sensors) magnitude of grand average regression coefficients (*β*). Sensor topographies for time points indicated by the black dots are shown below the main panel. Note that for the time points before the response we use a different scaling of colours than for time points around the response and later. This is to more clearly visualise the topography around the response which contains larger values. The colour scaling for the time points before the response is equal to that of Figure 3 and Figure 4. The topography at −300 ms strongly resembled that for accumulated evidence, but the topography around the response (30 ms) additionally exhibited stronger fronto-lateral and weaker occipital anti-correlations (*p* < 0.01 corrected, cf. Supplementary Figure 6). Positive values / correlations mean that measured sensor values tended to be high for a right choice (button press) and low for a left choice and vice-versa for negative values. See Supplementary Figure 5 to see how the central topography at 30 ms shown here results as the difference of the topographies associated with right and left choices.

The motor response itself (peak around 30 ms) was, as expected, much more strongly represented in the MEG signals than the accumulated evidence, see Figure 7. Although the motor response also had a predominantly central topography, its topography visibly differed from that prior to the response (at −300 and −120 ms). Specifically, the topography before the response exhibited stronger anti-correlation in occipital sensors than around the response while the topography around the response exhibited stronger anti-correlations in fronto-lateral sensors (*p* < 0.01 corrected, cf. Supplementary Figure 6). Furthermore, the correlation with choice was relatively higher over central sensors at 30 ms than at −120 ms (Supplementary Figure 6).

To analyse this difference at the source level we applied the regression analysis to the reconstructed source currents. Figure 8 depicts the results of an analysis of two time windows: the “build-up” window from −500 ms to −120 ms (when a dip before the response indicates an end of the build-up) and the “re-sponse” window capturing the response peak from −30 ms to 100 ms. We only show brain areas with at least one significant effect within the time window after correcting for multiple comparisons (FDR with *α* = 0.01 across brain areas and the two time windows). The shown colours indicate normalised second-level t-value magnitudes (see Methods).

**Figure 8:**
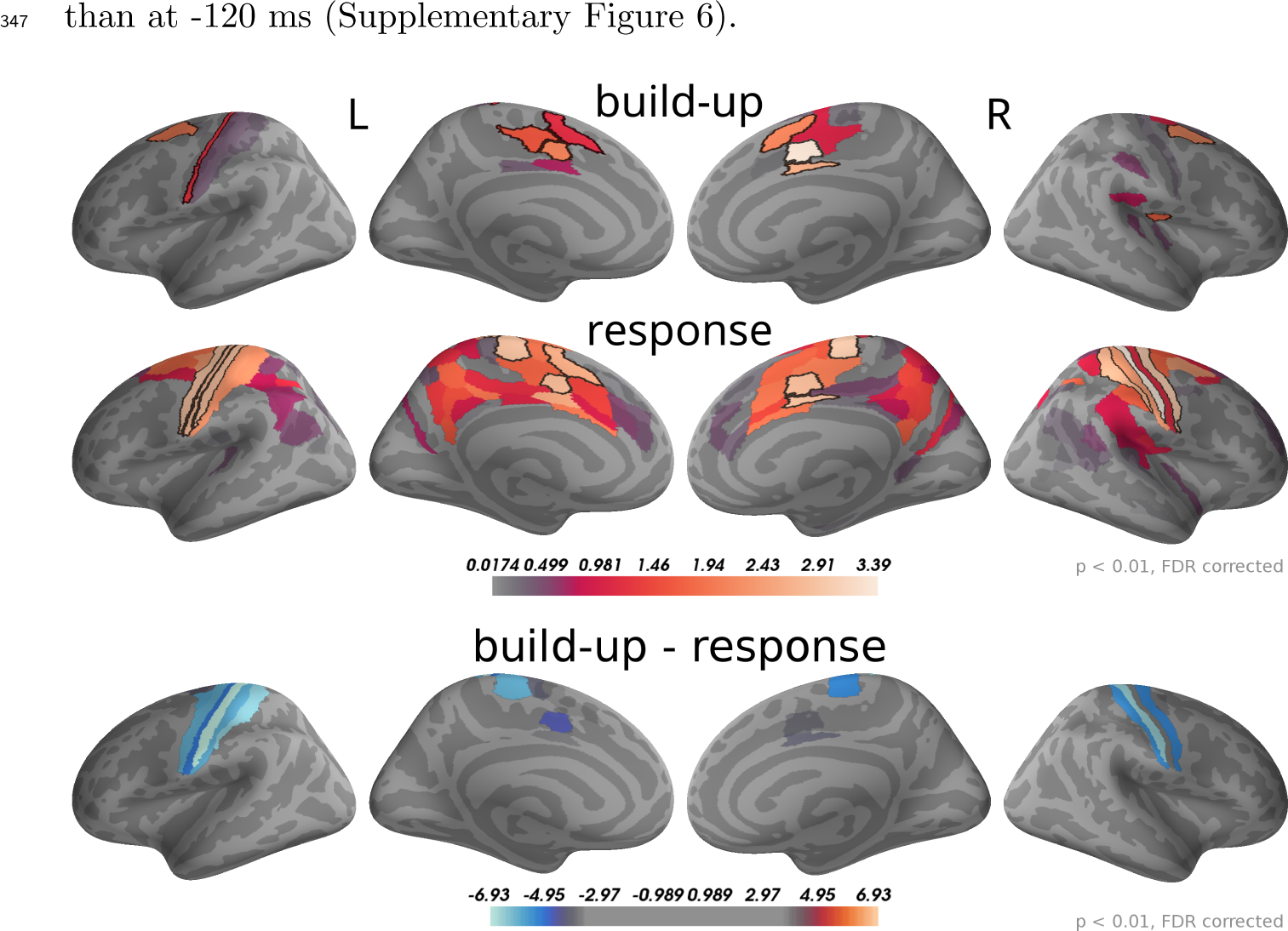
Around the response time strongest correlations with choice occurred in primary motor, somatosensory and cingulate motor cortex (BA 24) while during the build-up period we found the strongest effects in premotor and cingulate motor cortex. The 5 largest effects per hemisphere were: build-up, left – 24dv, 6a, 24dd, 3a, SCEF; right – 24dv, p24pr, 6a, SCEF, OP2-3; response, left – 4, 3b, 24dv, 3a, SCEF; right – 3b, 4, p24pr, 24dv, 2. When testing for differences in the spatial pattern of correlation magnitudes (see Methods) between the two time windows, we only found significant differences in the motor and cingulate areas: 1, 24dv, 2, 31a, 3a, 3b, 4, 6d, 6mp, SCEF, p24pr. All of these effects indicated that correlations with choice were stronger in the response window (blue). The build-up and response panels show spatially normalised t-value magnitudes while the difference panel shows t-values of spatially normalised correlation magnitude differences.

As expected, in the response window, the effects were dominated by choice correlations in bilateral primary motor and somatosensory cortices, but also choice correlations in cingulate motor areas (around Brodmann area 24) were among the effects with the strongest magnitudes. Other significant correlations with choice within the response window occurred in premotor and posterior cingulate cortices. In the build-up window, the strongest correlations occurred predominantly in cingulate motor cortex and premotor areas (especially 6a).

We further aimed at identifying brain areas with significantly different correlation magnitudes in the two time windows. Specifically, we were interested in the difference of the spatial patterns of correlation magnitudes, across brain areas, between the two time windows. To do this, we normalised correlation magnitudes across brain areas within the time windows and computed the differences between time windows within each brain area and participant (see Methods for details). Figure 8 bottom panel, shows that across participants the only statistically significant differences occurred in the primary motor and somatosensory cortices and, with smaller effect size, in cingulate motor areas. In all these areas correlation magnitudes were larger in the response as compared to the build-up window.

In summary, the response-centred analysis of choice correlations suggests that the build-up of choice-correlations leading towards a response is related to the accumulation of momentary evidence, because sensor topographies and brain areas were highly consistent across choice- and evidence-based analyses. The correlation topographies for the build-up and the response windows shown in Figure 7 had significant differences in central, occipital and fronto-lateral sensors. When analysing these differences at the source level (Figure 8), the only sources with significant differences were located in motor areas. These results together suggest that the brain areas representing decision evidence are largely overlapping with those representing the upcoming choice and the motor response. The difference in correlation patterns at the source level between the upcoming choice and motor response could be explained by an increase in choice correlations in motor areas.

## 3 Discussion

Using MEG, we have analysed the dynamics of evidence representations in the human brain during perceptual decision making. We induced fast, within-trial evidence fluctuations using a visual stimulus in which new, momentary evidence appeared every 100 ms and correlated the resulting momentary evidence dynamics with MEG signals. We found that each update of momentary evidence elicited a stereotyped response in the MEG signal that lasted until about 600 ms after the update onset, meaning that the brain processed incoming pieces of momentary evidence in parallel. We identified three main phases of the representation of momentary evidence: an early phase around 120 ms after an evidence update, an intermediate phase around 180 ms and a late phase from about 300 to 500 ms. These phases exhibited different sensor topographies with positive correlations shifting from occipital to centro-parietal to central sensors during the three phases. Using source reconstruction, we localised these representations of momentary evidence in early visual, parietal and motor areas, respectively, with significant correlations in posterior cingulate cortex occurring in all three phases. Significant correlations with accumulated evidence including the most recent evidence update occurred continuously until about 550 ms after update onset and exhibited a central topography similar to that in the late phase of momentary evidence representations with corresponding sources. Additionally, response-aligned correlations of the MEG signal with the final choice of the participants shared a similar topography in a build-up phase hundreds of milliseconds before the response. The correlation analysis at the source level further showed that the only significant differences between build-up phase and motor response were higher choice correlations in motor areas during the response.

These results consolidate a wide range of separate previous findings: It has previously been shown that the human brain elicits electromagnetic signals that correlate with individual pieces of momentary evidence (Wyart et al., 2012; de Lange et al., 2010; Gluth et al., 2013; Gould et al., 2012). Compared to these studies we here for the first time used a reaction time paradigm with fast evidence changes every 100 ms, more directly mimicking natural perceptual decision making processes. More importantly, our results are the first to track momentary evidence representations through the three phases that we identified and the corresponding areas in the human brain, although at least the early and late phases were previously hinted at (Wyart et al., 2012).

A large proportion of previous work investigating the dynamics of evidence representations in the human brain focused on oscillatory signals (Donner et al., 2009; de Lange et al., 2013; Gould et al., 2012; Siegel, Donner, Oostenveld, Fries, & Engel, 2007). For example, it has been found that the average amount of evidence in a trial is represented in the power of oscillations in occipital and parietal cortex (Siegel et al., 2007). Further, the difference in the power of oscillations between central-left and central-right sensors exhibits an evidence-dependent build-up towards the response that appears to be generated in motor areas (Donner et al., 2009; de Lange et al., 2013). We here made corresponding observations, but directly in the trial-wise temporal MEG signals reflecting trial-wise signal variations correlated with decision evidence that are believed to result from minute, event-related fluctuations in the voltage potentials of neuronal populations.

There is overwhelming evidence that motor areas including areas in the premotor and primary motor cortex are involved in perceptual decision making, e.g. (Hanks & Summerfield, 2017; Heekeren, Marrett, & Ungerleider, 2008). Specifically, it has been shown that some single neurons in primary motor cortex represent momentary evidence (Thura & Cisek, 2014), that the strength of muscle reflex gains is proportional to the average amount of momentary evidence within a trial (Selen et al., 2012), that motor-evoked potentials can be related to accumulated evidence (Michelet et al., 2010; Hadar, Rowe, Di Costa, Jones, & Yarrow, 2016) and that classical lateralised readiness potentials which are thought to represent motor processes (Smulders & Miller, 2012) also exhibit evidence-dependent build-up in a detection task (Kelly & O’Connell, 2013). Our results further substantiate these findings by showing that human motor areas represent each update of momentary evidence roughly within 300 to 500 ms after the update onset and that accumulated evidence is represented in motor areas throughout the decision making process. Using a response-aligned analysis of choice-dependent effects in the same reference frame as the analyses of evidence, we could further show that the stimulus-aligned evidence representations resemble closely the representation of the final choice during a build-up phase before the motor response. This supports the hypothesis that previous observations of pre-response representations of an upcoming choice, such as the lateralised readiness potential, should be interpreted as expressions of an ongoing decision making process about the next sensible motor response. In sum, the present and previous findings strongly affirm a tight coupling between decision making and motor processes, as, for example, formulated in the affordance competition hypothesis (Cisek & Kalaska, 2010; Cisek, 2007), but also other theories in cognitive computational neuroscience (O’Regan & Noe, 2001; Clark, 2013; Friston, Daunizeau, & Kiebel, 2009).

One potential caveat of our correlation results in motor areas is that due to the specifics of our task participants may actually have executed micromovements trying to track the changes of the perceptual stimulus either with their eyes, or with minimal finger movements close to the response buttons. In this scenario the observed correlations in motor areas would be possibly explained by motor signals to the muscles. Although we cannot completely exclude this possibility we deem it unlikely, because: i) Stimuli were shown only very centrally at visual angles within about 10° visual angle with most stimuli within 5° diameter from fixation meaning that most of them were well within the foveal visual field. ii) The sensor topographies representing evidence were very similar to that associated with the motor response, that is, the evidence representations do not appear to be specifically related to eye movements. iii) As mentioned above, a large body of work employing a wide variety of different tasks already supports the reverse interpretation that motor areas represent decision evidence before motor execution. In conclusion, we do not believe that the correlations with momentary or accumulated evidence observed in motor areas of the brain are merely an expression of motor control signals that caused stimulus-correlated micro-movements. Even if such micro-movements existed, we deem it likely that these follow the time-course of decision evidence rather than decision-irrelevant stimulus properties, as suggested by recent results about the adaptation of reflex gains and motor evoked potentials during decision making (Selen et al., 2012; Michelet et al., 2010; Hadar et al., 2016).

The early, intermediate and late phases of momentary evidence representations mirrored the presumed general transfer of behaviourally relevant visual information through the brain (Kandel, Jessell, Schwartz, Siegelbaum, & Hudspeth, 2012). In the early phase around 120 ms after evidence updates we found the strongest representations of momentary evidence in early visual cortex and occipito-temporal areas while in the intermediate phase around 180 ms momentary evidence representations included areas in inferior and superior parietal cortex. In the late phase the momentary evidence was predominantly represented in pre-/motor, somatosensory and cingulate areas while we only found one weak significant correlation with momentary evidence in one area of parietal cortex (right PFt). The same was true for representations of accumulated evidence. Taken together, these results suggest that in our task parietal cortex was only transiently involved in the processing of momentary evidence and that it did not accumulate evidence for the decision, or at least did not represent accumulated evidence over an extended period of time.

These results appear to be at odds with previous findings in non-human primates which had identified neurons in inferior parietal cortex that seemed to represent accumulated evidence (Gold & Shadlen, 2007). More recent work, however, suggests that the firing of these neurons is more diverse than originally thought (Latimer, Yates, Meister, Huk, & Pillow, 2015; I. M. Park, Meister, Huk, & Pillow, 2014; Meister, Hennig, & Huk, 2013). It is possible that the signal from only few evidence accumulating neurons in inferior parietal cortex is too weak to be recorded with MEG. Another possibility why we do not find strong correlations with accumulated evidence in parietal areas is that probably these representations are not as strongly lateralised in parietal areas as they are in motor areas. This would make it much harder to detect them with the typically low spatial resolution of MEG. Yet another possibility is that the representation of decision evidence in parietal areas follows a more intricate dynamic process that is hard to identify with simple correlation analyses (Churchland et al., 2010). If this was the case, an interesting follow-up question would be why the representations of accumulated evidence in parietal and motor or posterior cingulate areas apparently differ, as we clearly found correlations with accumulated evidence in the latter areas.

To manipulate decision evidence in our task we changed the position of a single dot presented on a screen. Only the x-coordinates of these dot positions represented momentary decision evidence while the decision-irrelevant y-coordinates acted as a perceptual control variable. We have shown that correlations of MEG signals with the perceptual control variable, in contrast to momentary evidence, were strongly diminished in the period from 300 to 500 ms after dot onset. This suggests that the brain ceases to represent perceptual information that is behaviourally irrelevant around this time and that brain areas with strong correlations with momentary evidence in this time window indeed are involved in the decision making process. This interpretation is further supported by previous work which has shown that purely perceptual stimulus information is represented in electrophysiological signals only until about 400 ms after stimulus onset (Wyart et al., 2012; Myers et al., 2015; Mostert, Kok, & de Lange, 2015) while specifically decision-related information is represented longer starting around 170 ms after stimulus onset (Wyart et al., 2012; Myers et al., 2015; Mostert et al., 2015; Philiastides & Sajda, 2006; Philiastides, Ratcliff, & Sajda, 2006; Philiastides, Heekeren, & Sajda, 2014).

We further validated this interpretation by investigating correlations with accumulated evidence, that is, the cumulative sum of momentary evidences within a trial. In contrast to the momentary evidence, this sum is more specifically related to the decision and has no simple, purely perceptual interpretation. The similarity of the topographies for accumulated evidence correlations and for momentary evidence correlations in the late phase suggests that specifically decision-relevant evidence is represented in the late phase, that is, within 300 to 500 ms after evidence updates. Our results do not allow to clearly state whether momentary, or accumulated, or both types of decision evidence were represented in the brain in this time window, because both types of evidence are correlated, especially early within a trial. However, we also found that accumulated evidence exhibited the corresponding central topography more consistently throughout peri-stimulus time than momentary evidence, so it appears reasonable to assume that predominantly accumulated evidence is represented in the late phase.

Finally, and perhaps most surprisingly, we found significant correlations with momentary and accumulated evidence in posterior cingulate cortex across all the investigated phases. Especially a ventral part of posterior cingulate cortex (v23ab) was involved already in the early phase which was dominated by correlations of momentary evidence in early visual areas and may therefore relate to basic visual processing of the stimulus. In the intermediate phase, the correlations in posterior cingulate cortex were weaker, but persisted. In the late phase correlations in posterior cingulate cortex constituted one of the main effects suggesting that it is a region contributing to the maintenance and accumulation of momentary evidence in the brain. Consequently, posterior cingulate cortex appears to be involved in both early sensory processing and decision making and, therefore, could act as a bridge between these processes.

Previous studies investigating the function of posterior cingulate cortex have mostly concentrated on a rather slow time scale, for example, contrasting different task conditions to each other, while we analysed rapid fluctuations of neural signals. These studies of slow changes in posterior cingulate cortex activations have implicated the posterior cingulate as having a direct role in directing the focus of attention (Leech & Sharp, 2014). However, posterior cingulate cortex has been associated with a wide range of functions which have recently been summarized as estimating the need to change behaviour in light of new, external requirements (Pearson, Heilbronner, Barack, Hayden, & Platt, 2011). Our findings are compatible with this view, when transferred to the context of comparably fast perceptual decision making where decision evidence may be viewed as the need to follow one (press left) or another (press right) behaviour.

In the field of perceptual decision making, especially in electrophysiological work with non-human animals, the posterior cingulate cortex has not gained much attention (Gold & Shadlen, 2007; Hanks & Summerfield, 2017). Given our findings it, therefore, appears that the role of posterior cingulate cortex in perceptual decision making may have been underestimated.

## 4 Materials and Methods

This study has been approved by the ethics committee of the Technical University of Dresden (EK324082016). Written informed consent was obtained from all participants. Code implementing the statistical analysis which produced all presented results is available at https://github.com/sbitzer/BeeMEG.

### 4.1 Participants

37 healthy, right-handed participants were recruited from the Max Planck Institute for Human Cognitive and Brain Sciences (Leipzig, Germany) participant pool (age range: 20 – 35 years, mean 25.7 years, 19 females). All had normal or corrected-to-normal vision, and reported no history of neurologic or psychiatric disorders. One participant was excluded from MEG measurement due to low performance during training. In total, 36 participants participated in the MEG study. Two participants’ data were excluded from analyses due to excessive eye artefacts and too many bad channels. Finally, 34 participants’ data were analysed (age range: 20 – 35 years, mean 25.85 years, 17 females).

### 4.2 Stimuli

In each trial, a sequence of up to 25 white dots were presented on a black screen. Each dot was displayed for 100 ms (6 frames, refresh rate 60 Hz). The white dot was located at x, y coordinates which were sampled from one of two twodimensional Gaussian distributions with means located at ±25 pixels horizontal distance from the centre of the screen. The standard deviation was 70 pixels in both axes of the screen. The mean locations were the two target locations (−25: left, 25: right). These target locations corresponded to visual angles ±0.6° from the centre of the screen. The standard deviation of the Gaussian distribution corresponded to ±1.7° from the two target locations. The stimuli used in this study consisted of a subset of stimuli used previously(H. Park et al., 2016), and additional newly created stimuli. The stimuli were chosen to increase the probability that the participants see the 5th dot within the 25 dot sequence by not responding earlier. In short, trials where 70% of the participants in the previous study (H. Park et al., 2016) had reaction times (RT) longer than 700 ms but not timed-out were chosen from the second most difficult condition. This resulted in 28 trials from 200 trials. Then each trial was copied 6 times, with only the 5th dot location differing, ranging in ‘target location + [−160 -96 −32 32 96 160] (pixels)’. This resulted in 168 trials. These trials were mirrored to create a dataset with the same evidence strengths but with different x coordinate signs (336 trials), and finally trials which had short RTs were chosen from (H. Park et al., 2016) as catch trials, to prevent participants from adapting to the long RT trials (30% of the total trials). This resulted in a total of 480 trials per experiment.

We originally designed this stimulus set, especially the manipulations of the 5th dot, to increase the chance of inducing sufficiently large effects in the MEG signal when observing the 5th dot. In a preliminary analysis we realised, however, that the natural variation of the stimuli already induces observable effects. Consequently, we pooled all trials for analysis.

### 4.3 Procedure

Participants were seated in a dimly lit shielding room during the training and the MEG measurement. Visual stimuli were presented using Presentation® software (Version 16.0, Neurobehavioral Systems, Inc., Berkeley, CA, www.neurobs.com). The display was a semi-transparent screen onto which the stimuli were back-projected from a projector located outside of the magnetic shielding room (Vacuumschmelze Hanau, Germany). The display was located 90 cm from the participants. The task was to find out which target (left or right) was the centre of the white dot positions, but participants were instructed with a cover story: Each target represented a bee hive and the white dot represented a bee. Participants should tell which bee hive is more likely the home of the bee. They were additionally instructed to be both accurate and fast, but not too fast at the expense of being inaccurate, and not too slow that the trial times out. They went through a minimum 210 and maximum 450 trials of training, until they reached a minimum of 75% accuracy. Feedback (correct, incorrect, too slow, too fast) was provided during the training. After training, a pseudo-main block with 200 trials without feedback preceded MEG measurement. After the pseudo-main session, the 480 trials in randomized order were presented to each participant divided into 5 blocks. The MEG measurement lasted 60 minutes, including breaks between blocks. Each trial started with a fixation cross (randomized, 1200 ms 1500 ms uniform distribution) followed by two yellow target dots. After 700 ms, the fixation cross disappeared and the first white dot appeared. The white dot jumped around the screen and stayed at each location for 100 ms, until the participant submitted a response by pressing a button using either hand, corresponding to the left / right target, or when the trial timed-out (2.5 s). In order to maintain motivation and attention throughout the measurement, participants were told to accumulate points (not shown to the participants) for correct trials and adequate (not too slow and not too fast, non-time-out) RTs. Bonus money in addition to compensation for participating in the experiment were given to participants with good performances. RTs and choices were collected for each trial for each participant. Although the trial order was randomized across participants, every participant saw exactly the same 480 trials.

### 4.4 Model of decision making behaviour

We used a previously described ideal observer model of decision making behaviour that is equivalent to a drift-diffusion model to define decision evidence (H. Park et al., 2016; Bitzer et al., 2014). The model postulates a direct linear relationship between momentary decision evidence and the x-coordinates of the white dot and identifies accumulated evidence as the simple cumulative sum of x-coordinates. Parameters of the model, that are typically fit to behavioural responses, only change the slope, or offset of the linear relationship between x-coordinates and momentary decision evidence. As we normalised x-coordinates before entering them in subsequent analyses, these parameters of the model are irrelevant for our purposes. Therefore, the decision making model had no further role in our analyses than providing the theoretical link between x-coordinates and momentary and accumulated decision evidence.

### 4.5 MEG data acquisition and preprocessing

MEG data were recorded with a 306 channel VectorviewTM device (Elekta Oy, Helsinki, Finland), sampled at 1000 Hz. The MEG sensors covered the whole head, with triplet sensors consisting of two orthogonal gradiometers and one magnetometer at 102 locations. Additionally, three electrode pairs were used to monitor eye movement and heart beats at the same sampling rate. The raw MEG data was corrected for head movements and external interferences by the Signal Space Separation (SSS) method (Taulu, Simola, & Kajola, 2005) implemented in the MaxFilterTM software (Elekta Oy) for each block. The subsequent preprocessing was performed using MATLAB (Mathworks, Massachusetts, United States). The head movement corrected data was high-pass and low-pass filtered using a linear phase FIR Kaiser filter (corrected for the shift) at cut-off frequencies of 0.33 Hz and 45 Hz respectively, with filter orders of 3736 and 392, respectively. The filtered data was then down-sampled to 250 Hz. Then independent component analysis (ICA) was applied to the continuous data using functions in the EEGLAB (Delorme & Makeig, 2004) to remove eye and heart beat artefacts. The data dimensionality was reduced by principal component analysis to 50 or 60 components prior to running the ICA. Components which had high temporal correlations (> 0.3) or typical topographies with/of the EOG and ECG signals were identified and excluded. The ICA-reconstructed data for each block was combined, and epoched from - 300 ms to 2500 ms from the first dot onset (zero). Another ICA was applied to these epoched data in order to check for additional artefacts and confirm typical neural topographies from the components. The ICA reconstructed data and original data were compared and inspected in order to ensure only artefactual trials were excluded. Before statistical analysis we used MNE-Python v0.15.2 (Gramfort et al., 2014, 2013) to downsample the data to 100 Hz (10 ms steps) and perform baseline correction for each trial where the baseline value was the mean signal in the period from −300 ms to 0 ms (first dot onset).

### 4.6 Source reconstruction

We reconstructed the source currents underlying the measured MEG signals using noise-normalised minimum norm estimation (Dale et al., 2000) implemented in the MNE software. To create participant-specific forward models we semi-automatically co-registered the head positions of participants with the MEG coordinate frame while at the same time morphing the participants’ head shape to that of Freesurfer’s fsaverage by aligning the fsaverage head surface to a set of head points recorded for each participant. We defined a source space along the white matter surface of the average subject with 4098 equally spaced sources per hemisphere and an approximate source spacing of about 5 mm (MNE’s “oct6” option). For minimum norm estimation we assumed a signal-to-noise ratio of 3 (lambda2 = 0.11). We estimated the noise covariance matrix for noise normalisation (Dale et al., 2000) from the MEG signals in the baseline period spanning from 300 ms before to first dot onset in each trial. We further used standard loose orientation constraints (loose=0.2), but subsequently picked only the currents normal to the cortical mantle. We employed standard depth weighting with a value of 0.8 to overcome the bias of minimum norm estimates towards superficial sources. We computed the inverse solution from all MEG sensors (magnetometers and the two sets of gradiometers) returning dynamic statistical parametric maps for each participant. Before some of the subsequent statistical analyses we averaged the reconstructed source signals across all sources of a brain area as defined by the recently published HCP-MMP parcellation of the human connectome project (Glasser et al., 2016).

### 4.7 Regression analyses

Most of our results were based on regression analyses with a general linear model giving event-related regression coefficients (Clarke et al., 2013; Hauk et al., 2006). We differentiate between a standard regression analysis on events aligned at the time when the white dot appeared in each trial, expanded regression analyses on events aligned at the times of white dot position changes and response-aligned regression analyses.

#### 4.7.1 Standard regression analysis

In the standard regression analysis we defined dot-specific regressors with values changing only across trials. For example, we defined a regressor for momentary evidence (x-coordinate) of the 2nd white dot position presented in the trial. For convenience we also call white dot positions (1st, 2nd and so forth in the sequence of dot positions) simply ‘dots’.

We only report results of a standard regression analysis in Supplementary Figure 1. This analysis included the dot x- and y-coordinates of the first 6 dots as regressors of interest (together 12 regressors). Additional nuisance regressors were: the response of the participant, a participant-specific trial count roughly measuring time within the experiment, an intercept capturing average effects and a response entropy. The latter quantified the posterior uncertainty of a probabilistic model of the responses (H. Park et al., 2016) that the model had about the response for the stimulus presented in that trial after model parameters were adapted to fit participant responses. Specifically, the wider and flatter the posterior predictive distribution over responses of the model for a particular trial / dot position sequence was, the larger was the response entropy for that trial. The data for this analysis were the preprocessed magnetometer time courses.

#### 4.7.2 Expanded regression analyses

Expanded regression analyses were based on an expanded set of data created by dividing up the data into partially overlapping epochs centred on the times of dot position changes. For each time point after this dot onset the data contained a variable number of time points depending on how many more dots were presented in each individual trial before a response was given by the participant. For example, if a participant made a response after 880 ms in a trial, 9 dots were shown in that trial (onset of the 9th dot was at 800 ms). If we are interested in the time point 120 ms after dot onset (dot position change), this gives us 8 time points within that trial that were 120 ms after dot onset. Further excluding all time points 200 ms before the response and later, would leave us with 6 data points for this example trial. See Figure 9 for an illustration. For each time after dot onset and for each participant we pooled all of these data points across trials and inferred regression coefficients on these expanded data sets. Note that this approach can equally be interpreted as statistical inference over how strongly the sequence of momentary evidence caused by the dot updates is represented in the signal at 100 ms wide steps with a delay given by the chosen time from dot onset.

**Figure 9:**
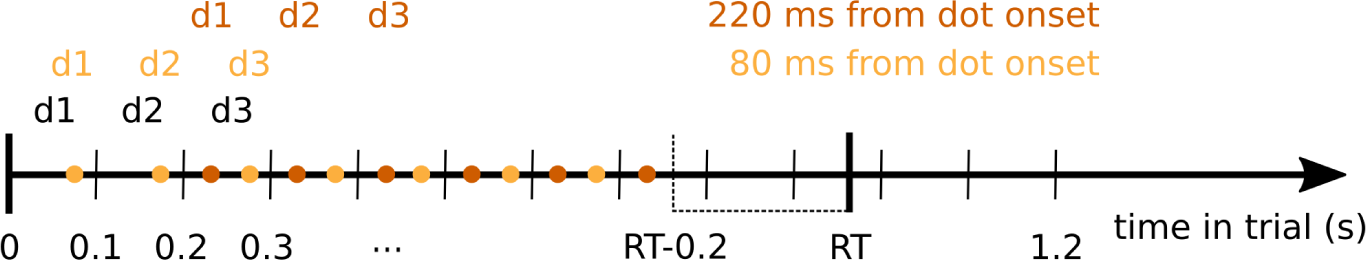
Diagram demonstrating the selection of data points entering the expanded regression analyses. Dot positions (d1, d2, d3, …) changed every 100 ms in the experiment (black). Coloured dots indicate times at which signal data points entered the analysis for a given time from dot position change (dot onset, shown exemplarily for 80 and 220 ms from dot onset). We only considered time points up to 200 ms before the response in each trial. Coloured d1, d2, d3 above the points indicate the dot positions associated with the corresponding signal data points for the given time from dot onset. For each trial, these pairs of signal data and dot positions entered the expanded regression analyses.

These analyses included two regressors of interest: momentary evidence (x-coordinate) and y-coordinate of the associated dots. We additionally included the following nuisance regressors: an intercept capturing average effects, the absolute values of x- and y-coordinates, perceptual update variables for x- and y-coordinates (Wyart et al., 2012) defined as the magnitude of the change from one dot position to another and accumulated values of x- and y-coordinates. Because we found that the accumulated values can be strongly correlated with the individual x- and y-coordinates (cf. Supplementary Figure 2), we only used accumulated values up to the previous dot in the regressor. For example, if a data point was associated with the x-coordinate of the 4th dot, the accumulated regressor would contain the sum of only the first three x-coordinates. This accumulated regressor is equal to the regressor resulting from Gram-Schmidt orthonormalisation of the full sum of x-coordinates with respect to the last shown x-coordinate. The accumulated evidence regressor was derived from the ideal observer model as the log posterior odds of the two alternatives, but this was almost 100% correlated with the simple sum of x-coordinates. The small differences between model-based accumulated evidence and sum of x-coordinates after normalisation resulted from a small participant-specific offset representing the overall bias of the participant towards one decision alternative. Note that we do not show any results for this (previous) accumulated evidence regressor.

In Figure 4, Figure 6, Supplementary Figure 3 and Supplementary Figure 4 we report results from separate expanded regression analyses in which we replaced the x-coordinate regressor with the sum of x-coordinates and dropped the previous accumulated evidence regressor. We did this, because the previous accumulated evidence regressor did not allow us to estimate effects of accumulated evidence for the first 100 ms after dot onset which is possible with the separate regression. We also did not see any benefits from using the previous accumulated evidence regressor in comparison to the simple sum of x-coordinates up to the current dot. Although the previous accumulated evidence regressor is in principle Gram-Schmidt orthogonalised with respect to the current, i.e., last presented x-coordinate and therefore provides independent information from the current x-coordinate, this is not the orthogonalisation that we are most interested in. Ideally we would want to orthogonalise with respect to any information about x-coordinates, i.e., momentary evidence including information contributed by the whole series of x-coordinates. So, while the previous accumulated evidence regressor is orthogonal to the current x-coordinate, it still correlates with the x-coordinates of previously presented dots. As accumulated evidence is just the sum of x-coordinates, this cannot be prevented so that momentary and accumulated evidence regressors will always partially capture overlapping effects. We still found it informative to present a separate analysis for accumulated evidence under the premise that the effects of the accumulated evidence regressor more strongly relate to accumulated evidence than momentary evidence and vice-versa for the momentary evidence regressor. We present a discussion of their differences in Supplementary Material.

#### 4.7.3 Response-aligned regression analyses

Additional to the first-dot onset and dot onset aligned analyses, we conducted response-aligned analyses in which time was referenced to trial-specific response times of participants. The regressors in this analysis were the trial-specific choice of the participant, trial-time and an intercept. Choice was encoded as −1 for left and +1 for right so that the direction of correlations was compatible with that for the evidence regressors. The trial-time regressor simply counted the trial number within the experiment per participant. Timed out trials were excluded from analysis. As in the other regression analyses we z-scored regressors and data across trials before estimating the regression coefficients, except for trial-time which was only scaled to standard deviation equal to 1. We ran two different analyses in sensor and source space. In sensor space (magnetometers) we ran independent univariate regressions for each combination of sensor and time so that we ran 102 * 70 regressions with maximally 480 data points (one per trial, minus excluded trials). We report results of this analysis in Figure 7 and Supplementary Figure 6. After having identified time windows of interest based on the sensor level results, we aggregated data from the identified times into a common regression on source data. To do this we simply pooled the data from all times in the time window and ran the regression on this expanded data set, then including maximally number of trials * number of time points data points. This approach meant that we were automatically estimating the mean regression coefficients across the selected time window for each brain area and participant. We report results of this analysis in Figure 8.

#### 4.7.4 Identification of significant source-level effects

To identify significant correlations between regressors of interest and source signals we followed the summary statistics approach (Friston, Ashburner, Kiebel, Nichols, & Penny, 2006) and performed two-sided t-tests on the second level (group-level, t-tests across participants). We corrected for multiple comparisons across time points and brain areas by controlling the false discovery rate using the Benjamini-Hochberg procedure (Benjamini & Hochberg, 1995). Specifically, for identifying significant effects reported in Figure 5 we corrected across 25,340 tests covering 70 time points (0 to 690 ms from dot onset in 10 ms steps) and 362 brain areas (180 brain areas of interest per hemisphere plus one collection of sources per hemisphere that fell between the area definitions provided by the atlas). We report all significant effects of this analysis in Supplementary Data Table 1.

#### 4.7.5 Identification of significant differences in correlation patterns

We formally investigated the differences in correlation patterns of the response-aligned analysis between the two time windows of interest (Figure 8 Supplementary Figure 6). As we were interested in the differences between spatial patterns, we accounted for the overall increase in correlation magnitudes from build-up to response window by normalising the correlation magnitudes. This normalisation consisted of first shifting the minimum magnitude to 0 and then scaling the resulting magnitudes so that their mean equals 1 across sensors or brain areas. The initial shift of the magnitudes prevents excessive shrinking of magnitude variances for magnitude patterns with overall large magnitudes and ensures that the magnitudes have similar distributions across the involved sensors or brain areas in both considered time periods. We subsequently computed the differences between the selected time periods on the first level and report second-level (across participant) statistics.

The analysis on the source level in principle equalled that of the sensor level, but additionally accounted for the fact that most brain areas were not involved in encoding the choice. We achieved this by computing the normalisation parameters for a time window only across brain areas with a significant effect in this time window. However, we then computed magnitude differences for all brain areas with a significant effect in at least one of the time windows and proceeded with second-level statistics for these areas as before.

## 5 Acknowledgements

We would like to thank Yvonne Wolff-Rosier for helping with data acquisition.

## 6 Supplementary Material

### 6.1 Stereotyped temporal correlation profiles across evidence regressors

**Supplementary Figure 1:**
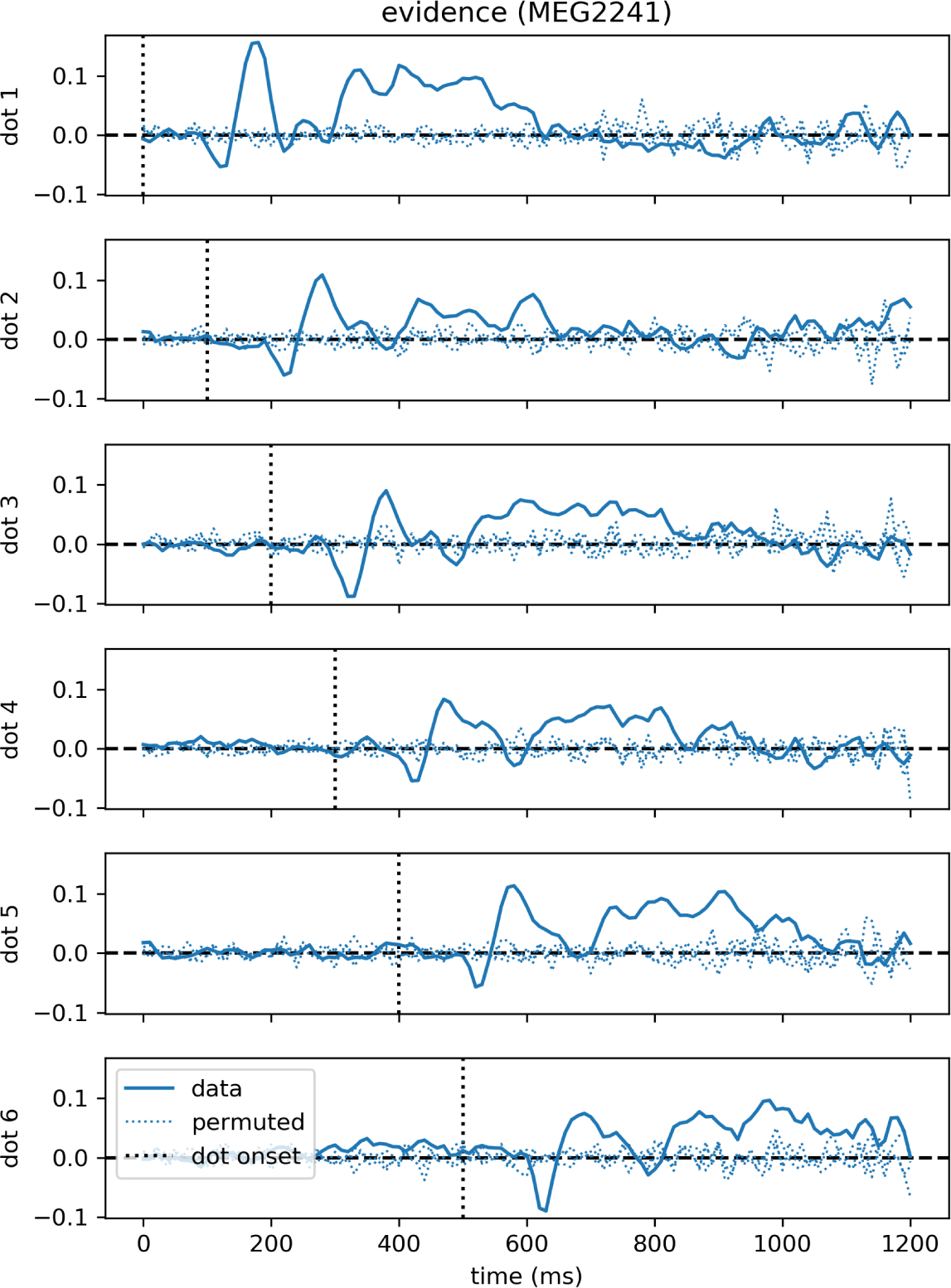
Time course of correlations with momentary evidence repeats for each dot shifted by dot onset times. In the standard regression analysis there was one regressor for each element in the sequence of dot positions (dots). This allowed us to see, when after first dot onset, correlations with the considered dot could be observed. The figure demonstrates exemplarily for the magnetometer channel with the strongest average correlations that the correlation time course exhibits roughly a stereotyped profile relative to the onset time of the dot on the second level. Dotted lines show the same quantity, but for data that we permuted over trials before the regression analysis.

### 6.2 Correlations with accumulated evidence

**Supplementary Figure 2:**
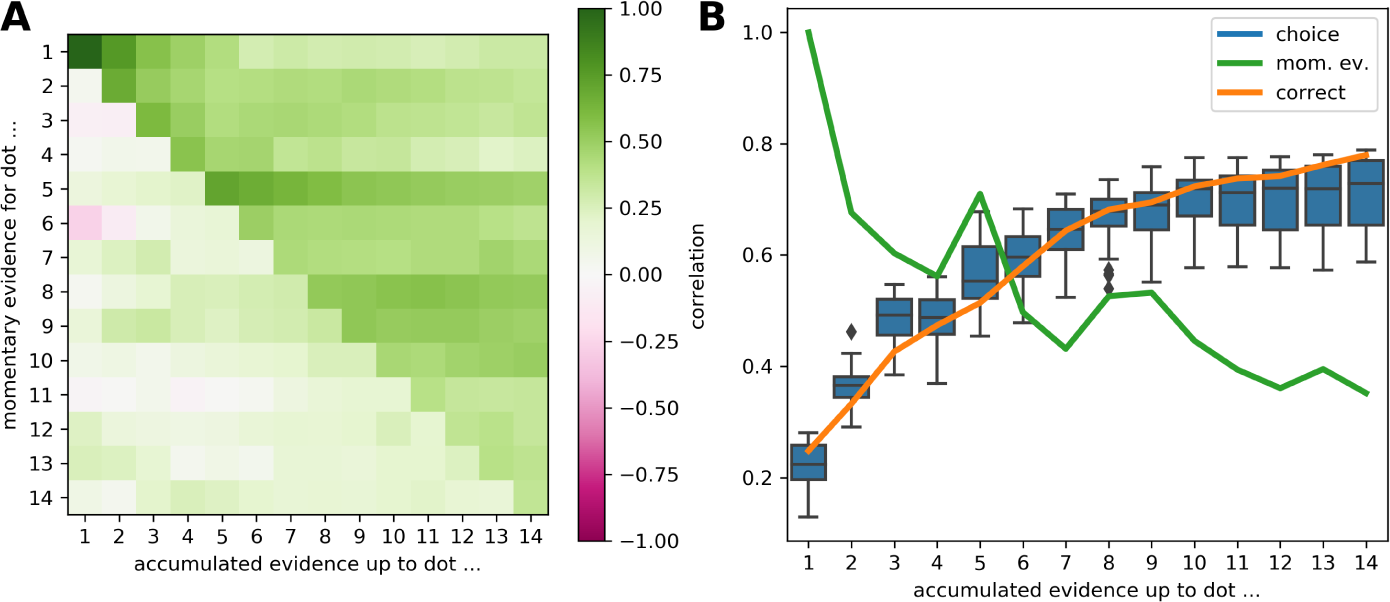
The accumulated evidence is correlated across trials with the momentary evidence provided by dot positions, the correct choice in a trial and the choices of the participants. A: Correlation coefficients for all combinations of momentary and accumulated evidence for the shown onset times. For example, the correlation value at row 2, column 4 gives the correlation between the momentary evidence of the 2nd dot position within a trial and the accumulated evidence up to the 4th dot position, across trials. B: Comparison of correlations between accumulated evidence and three trial-wise measures: the correct choice in a trial (orange line), the momentary evidence at the same time point (green line, equal to diagonal in A), and the choices of the participants (blue boxes). The blue boxes show the distribution over participants per considered dot position.

### 6.3 On the relation of momentary and accumulated evidence representations

The simple mathematical relationship between momentary and accumulated evidence (a sum) means that the inferred regression coefficients for either momentary or accumulated evidence will always contain contributions from the other type of evidence. In other words, a linear regression analysis (including any form of correlation analysis) will never be able to completely dissociate neural signals relating only to momentary or only to accumulated evidence. However, as inferred coefficients for momentary or accumulated evidence still identify effects of the corresponding type of evidence more strongly than the other type of evidence, we here attempt to further delineate effects relating to momentary and accumulated evidence by investigating specifically differences in their inferred representations. Our analysis suggests that specifically accumulated evidence and not momentary evidence is represented in the MEG magnetometer signals with a central positivity, as already indicated by the inferred coefficients for accumulated evidence shown in Figure 4.

Because accumulated evidence is the cumulative sum of momentary evidences, accumulated evidence can correlate strongly with the last shown momentary evidence and with momentary evidences shown at previous time points, even though momentary evidences at different time points are themselves uncorrelated. To understand the relation between momentary and accumulated evidence effects we, therefore, need to consider the recent history of momentary evidences.

To further visualise the relation between momentary and accumulated evidence we annotated the correlation time course of accumulated evidence shown in Figure 4 with that of the momentary evidence shown in Figure 3. This is shown in Supplementary Figure 3 where we additionally added two time-shifted replicas of the momentary evidence. These replicas visualise the representations of the x-coordinates of two previous dots such that we see the time points at which x-coordinates of the current and two previous dots are represented in the MEG signals while only the time course of correlations with accumulated evidence up to the current dot are shown. The figure shows that the location of peaks of accumulated evidence can be explained with the location of peaks of momentary evidence (x-coordinates). For example, at 180 ms peaks of accumulated evidence and momentary evidence of the current dot coincide. Similarly, the peak of accumulated evidence at 80 ms can be related to the 180 ms peak of momentary evidence of the previous dot (occurring at 80 ms in reference to the current dot onset time). These observations demonstrate that the correlations with accumulated evidence are partially driven by representations of the momentary evidence / x-coordinates in the MEG signals.

To disentangle representations of accumulated evidence and momentary evidence, or dot x-coordinates the time points with large differences in correlation strengths between the two are of greatest interest. The two time points with the largest discrepancies between correlation magnitudes of momentary and accumulated evidence were 20 ms and 120 ms after dot onset. Supplementary Figure 3 shows that at 20 and 120 ms a drop in magnitude of accumulated evidence correlations co-occurred with the 120 ms momentary evidence peaks of the current and previous dots. This drop in magnitude of accumulated evidence correlations, therefore, resulted from an interaction with the centro-parietal anti-correlation of the MEG signal with x-coordinates (cf. Figure 3A topography at 120 ms). Put differently, at these time points the representations of accumulated evidence and x-coordinates in the MEG signal were incompatible so that through the correlation between accumulated evidence and x-coordinates the correlations of accumulated evidence with the MEG signal were diminished as the MEG signal simultaneously represented the incompatible x-coordinates. Despite this interaction, however, we still observed positive correlations with accumulated evidence in central sensors mirroring the topography, although weaker, at later time points with strong effects, for example at 320 ms (Supplementary Figure 3 bottom). This affirms that specifically accumulated evidence is represented with a topography featuring a central positivity in the MEG signal.

**Supplementary Figure 3:**
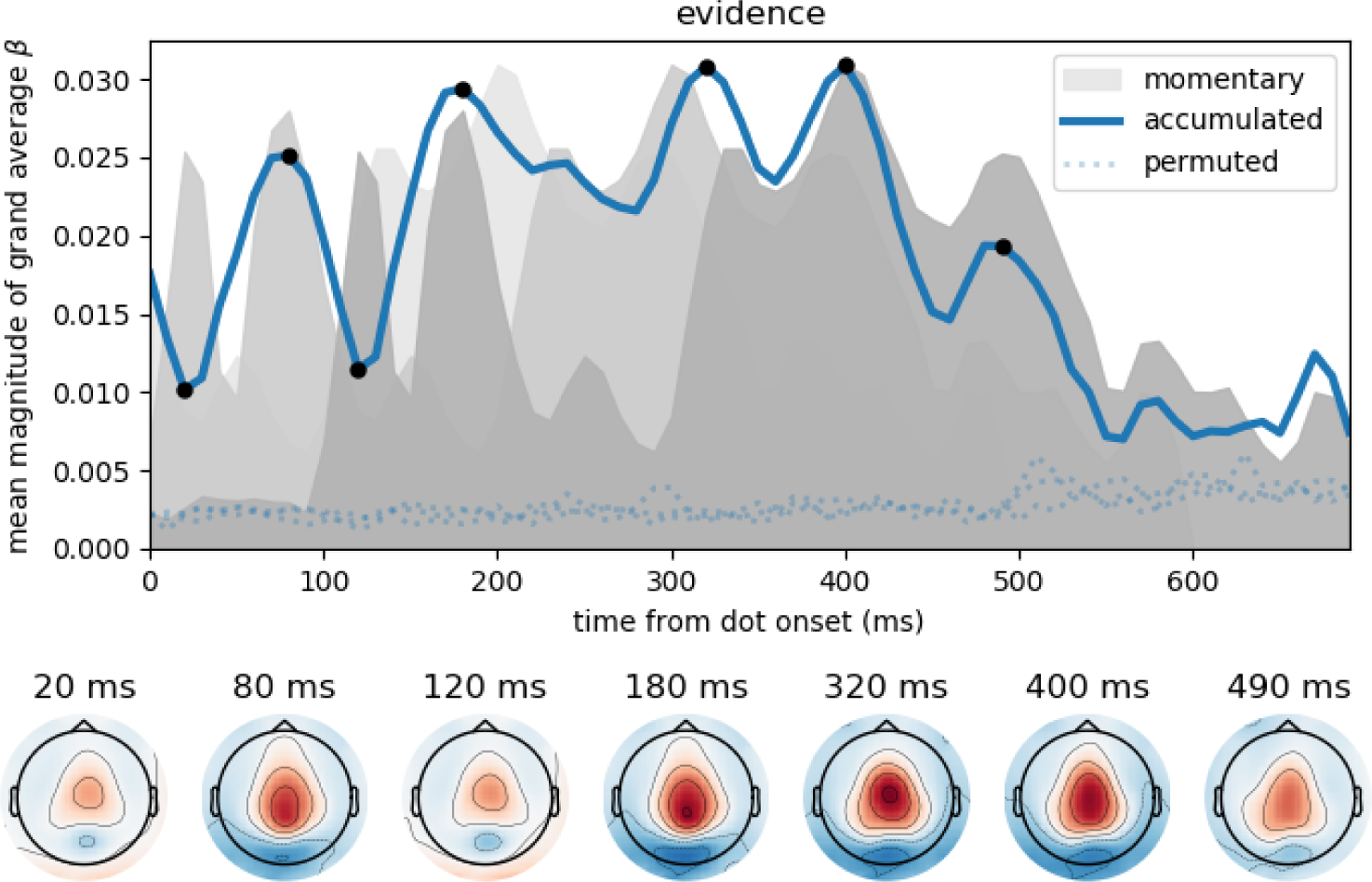
Peaks and troughs of accumulated evidence correlations coincide with peaks of momentary evidence correlations. Same format as in Figure 3. Accumulated evidence is plotted as a blue solid line, where black dots indicate time points of sensor topographies. In addition, we show the momentary evidence time course (dark grey shade, cf. Figure 3A) and two time-shifted replicas of it, one shifted by 100 ms to the past (mid grey) and another shifted by 200 ms to the past (light grey). These time courses, therefore, are associated with the representation of the momentary evidence / x-coordinates of the current and the previous two dots in the brain. This visualisation shows that peaks in accumulated evidence tend to coincide with peaks in momentary evidences presented at subsequent time points. Larger discrepancies between correlation magnitudes of momentary and accumulated evidence only occurred at 20 ms, 120 ms and from about 450 ms after dot onset. At 80 ms and 180 ms topographies slightly shifted towards parietal sensors otherwise effects were located centrally.

At 80 ms and 180 ms the sensor topographies for accumulated evidence deviated somewhat from a central to a more centro-parietal positivity. We reported above that the peaks of accumulated evidence coincided with peaks of momentary evidence at these time points. These momentary evidence peaks corresponded to the 180 ms momentary evidence peak in relation to dot onset which had a centro-parietal topography as shown in Figure 3A. So the shift from central to centro-parietal positive correlations with accumulated evidence can be explained by the interaction with the representation of the momentary evidence at these time points. Notice, however, that the positive correlations with accumulated evidence still cover central locations more strongly than the corresponding effects for momentary evidence at 180 ms (Supplementary Figure 4). This also indicates that accumulated evidence tended to be represented with a central positivity.

**Supplementary Figure 4:**
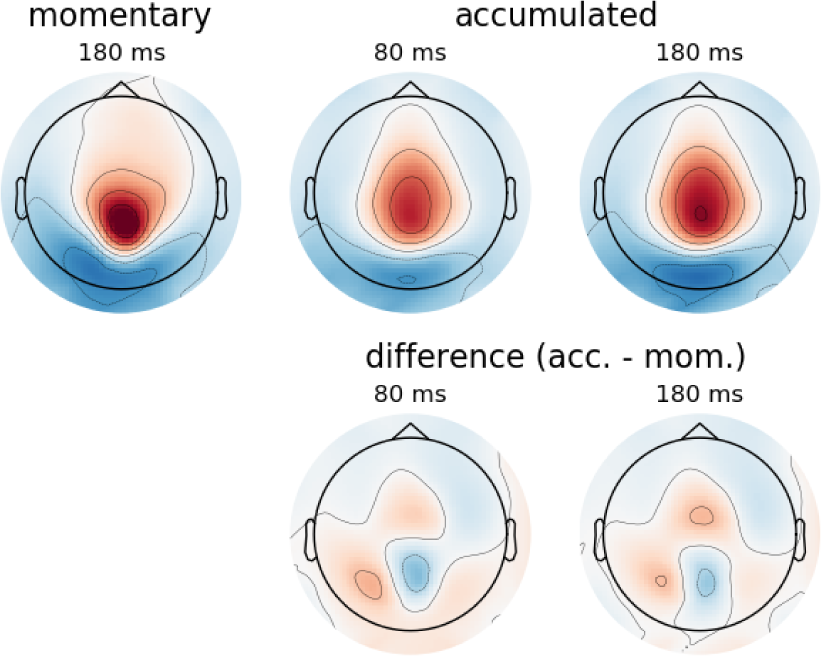
Correlations between magnetometer signals and accumulated evidence are more central than those for momentary evidence at 180 ms after dot onset. Top: Topographies repeated from Figure 3 and Supplementary Figure 3 at the indicated times. Bottom: Difference topographies where we subtracted the topography of the momentary evidence at 180 ms from those of the accumulated evidence. The difference topographies show that the accumulated evidence had stronger correlations in central sensors while the momentary evidence had stronger correlations in mid-parietal sensors. Additionally, accumulated evidence had stronger correlations in posterior lateral sensors, especially on the left.

### 6.4 Choice correlations correspond to difference between average signals for right minus left choices

**Supplementary Figure 5:**
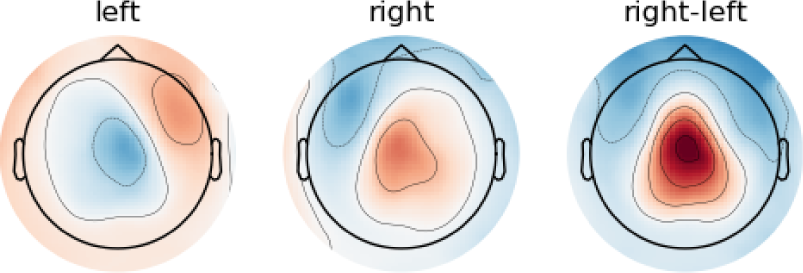
The choice correlations 30 ms after the response shown in Figure 7 emerge from the difference between right and left button responses. For this analysis we repeated the response-aligned regression analysis, but replaced the intercept and choice regressors with regressors for the left and right choices. The resulting regression coefficients are approximately proportional to the average signal in trials with a left, or right choice, respectively (the estimated coefficients are equal to the average signals, if the two response regressors are encoded with 0s and 1s and are the only regressors in the analysis, or their correlation with the other regressors is exactly 0). The topographies for left and right above show second-level t-values for the regression coefficients at 30 ms after the response. The topography on the right hand side shows their difference.

### 6.5 Topography differences for choice correlations before and around the response

**Supplementary Figure 6:**
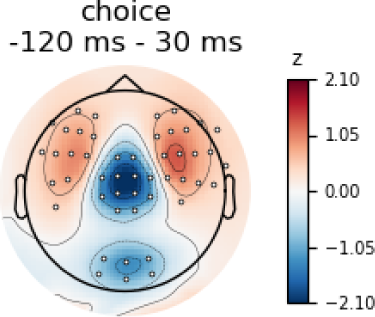
Around the response choice correlations were stronger than before the response over central sensors. We formally tested the apparent differences in topographies of choice correlations shown in Figure 7 for time points −120 ms and 30 ms. As we were interested in the spatial patterns and not absolute value differences within sensors, we scaled the coefficient estimates (*β*) across sensors, but within time points and participants for this analysis so that their mean magnitude across sensors was equal to 1. We then computed the difference between time points within each participant and sensor. The colouring in the plot shows the mean of these differences across participants. We further applied a t-test across participants within each sensor and corrected the resulting p-values for false discovery rate at *α =* 0.01 across sensors. The white dots in the figure indicate sensors which exhibited a significant difference after multiple comparison correction. Together with the topographies shown in Figure 7 the results of this analysis confirm that before the response occipital sensors had stronger anti-correlation with choice than around the response. In contrast, fronto-lateral sensors exhibited stronger anti-correlation around the response than before the response. Furthermore, the strongest difference occurred in central sensors which exhibited a relatively stronger correlation with choice around the response than before the response.

## References

Benjamini, Y., & Hochberg, Y. (1995). Controlling the false discovery rate: a practical and powerful approach to multiple testing. Journal of the Royal Statistical Society. Series B. Methodological, 57(1), 289–300.

Bitzer, S., Park, H., Blankenburg, F., & Kiebel, S. J. (2014). Perceptual decision making: Drift-diffusion model is equivalent to a bayesian model. Frontiers in Human Neuroscience, 8(102). doi:10.3389/fnhum.2014.00102

Brunton, B. W., Botvinick, M. M., & Brody, C. D. (2013, Apr). Rats and humans can optimally accumulate evidence for decision-making. Science, 340(6128), 95-98. doi:10.1126/science.1233912

Churchland, M. M., Yu, B. M., Cunningham, J. P., Sugrue, L. P., Cohen, M. R., Corrado, G. S.,… Shenoy, K. V. (2010, Mar). Stimulus onset quenches neural variability: a widespread cortical phenomenon. Nat Neurosci, 13(3), 369-378. Retrieved from http://dx.doi.org/10.1038/nn.2501 doi:10.1038/nn.2501

Cisek, P. (2007, September). Cortical mechanisms of action selection: the affordance competition hypothesis. Philosophical transactions of the Royal Society of London. Series B, Biological sciences, 362, 1585-1599. doi:10.1098/rstb.2007.2054

Cisek, P., & Kalaska, J. F. (2010). Neural mechanisms for interacting with a world full of action choices. Annu Rev Neurosci, 33, 269-298. doi:10.1146/annurev.neuro.051508.135409

Clark, A. (2013, Jun). Whatever next? predictive brains, situated agents, and the future of cognitive science. The Behavioral and brain sciences, 36, 181-204. doi:10.1017/S0140525X12000477

Clarke, A., Taylor, K. I., Devereux, B., Randall, B., & Tyler, L. K. (2013, January). From perception to conception: how meaningful objects are processed over time. Cerebral cortex (New York, N.Y. : 1991), 23, 187–197. doi:10.1093/cercor/bhs002

Dale, A. M., Liu, A. K., Fischl, B. R., Buckner, R. L., Belliveau, J. W., Lewine, J. D., & Halgren, E. (2000, April). Dynamic statistical parametric mapping: combining fmri and meg for high-resolution imaging of cortical activity. Neuron, 26, 55-67. doi:10.1016/S0896-6273(00)81138-1

de Lange, F. P., Rahnev, D. A., Donner, T. H., & Lau, H. (2013, Jan). Prestimulus oscillatory activity over motor cortex reflects perceptual expectations. J Neurosci, 33(4), 1400-1410. doi:10.1523/JNEUR0SCI.1094-12.2013

de Lange, F. P., Jensen, O., & Dehaene, S. (2010, Jan). Accumulation of evidence during sequential decision making: the importance of top-down factors. J Neurosci, 30(2), 731-738. doi:10.1523/JNEUR0SCI.4080-09. 2010

Delorme, A., & Makeig, S. (2004, March). Eeglab: an open source toolbox for analysis of single-trial eeg dynamics including independent component analysis. Journal of neuroscience methods, 134, 9-21. doi:10.1016/ j.jneumeth.2003.10.009

Donner, T. H., Siegel, M., Fries, P., & Engel, A. K. (2009, Sep). Buildup of choice-predictive activity in human motor cortex during perceptual decision making. Curr Biol, 19(18), 1581-1585. doi:10.1016/j.cub.2009.07. 066

Friston, K. J., Ashburner, J. T., Kiebel, S. J., Nichols, T. E., & Penny, W. D. (Eds.). (2006). Statistical parametric mapping: The analysis of functional brain images. Academic Press.

Friston, K. J., Daunizeau, J., & Kiebel, S. J. (2009). Reinforcement learning or active inference? PLoS One, 4 (7), e6421. doi:10.1371/journal.pone. 0006421

Glasser, M. F., Coalson, T. S., Robinson, E. C., Hacker, C. D., Harwell, J., Yacoub, E.,… Van Essen, D. C. (2016, August). A multi-modal parcellation of human cerebral cortex. Nature, 536, 171-178. doi:10.1038/nature18933

Gluth, S., Rieskamp, J., & Biichel, C. (2013, October). Classic eeg motor potentials track the emergence of value-based decisions. NeuroImage, 79, 394-403. doi:10.1016/j.neuroimage.2013.05.005

Gold, J. I., & Shadlen, M. N. (2007). The neural basis of decision making. Annu Rev Neurosci, 30, 535-574. doi:10.1146/annurev.neuro.29.051605. 113038

Gould, I. C., Nobre, A. C., Wyart, V., & Rushworth, M. F. S. (2012, Oct). Effects of decision variables and intraparietal stimulation on sensorimotor oscillatory activity in the human brain. J Neurosci, 32(40), 13805-13818. doi:10.1523/JNEUROSCI.2200-12.2012

Gramfort, A., Luessi, M., Larson, E., Engemann, D. A., Strohmeier, D., Brodbeck, C.,… Hamalainen, M. (2013, December). Meg and eeg data analysis with mne-python. Frontiers in neuroscience, 7, 267. doi:10.3389/fnins.2013.00267

Gramfort, A., Luessi, M., Larson, E., Engemann, D. A., Strohmeier, D., Brodbeck, C.,… Hämäläinen, M. S. (2014, February). Mne software for processing meg and eeg data. NeuroImage, 86, 446-460. doi:10.1016/j.neuroimage.2013.10.027

Hadar, A. A., Rowe, P., Di Costa, S., Jones, A., & Yarrow, K. (2016, November). Motor-evoked potentials reveal a motor-cortical readout of evidence accumulation for sensorimotor decisions. Psychophysiology, 53, 1721-1731. doi:10.1111/psyp.12737

Hanks, T. D., & Summerfield, C. (2017, January). Perceptual decision making in rodents, monkeys, and humans. Neuron, 93, 15-31. doi:10.1016/ j.neuron.2016.12.003

Hauk, O., Davis, M. H., Ford, M., Pulvermüller, F., & Marslen-Wilson, W. D. (2006, May). The time course of visual word recognition as revealed by linear regression analysis of erp data. NeuroImage, 30, 1383-1400. doi:10.1016/j.neuroimage.2005.11.048

Heekeren, H. R., Marrett, S., & Ungerleider, L. G. (2008, Jun). The neural systems that mediate human perceptual decision making. Nat Rev Neurosci, 9(6), 467-479. doi:10.1038/nrn2374

Hubert-Wallander, B., & Boynton, G. M. (2015). Not all summary statistics are made equal: Evidence from extracting summaries across time. Journal of vision, 15, 5. doi:10.1167/15.4.5

Kandel, E., Jessell, T., Schwartz, J., Siegelbaum, S., & Hudspeth, A. (2012). Principles of neural science, fifth edition. McGraw-Hill Education. Retrieved from http://www.principlesofneuralscience.com/

Kelly, S. P., & O’Connell, R. G. (2013, Dec). Internal and external influences on the rate of sensory evidence accumulation in the human brain. J Neurosci, 33(50), 19434-19441. doi:10.1523/JNEUROSCI.3355-13.2013

Latimer, K. W., Yates, J. L., Meister, M. L. R., Huk, A. C., & Pillow, J. W. (2015, Jul). Single-trial spike trains in parietal cortex reveal discrete steps during decision-making. Science, 349(6244), 184-187. doi:10.1126/science.aaa4056

Leech, R., & Sharp, D. J. (2014, January). The role of the posterior cingulate cortex in cognition and disease. Brain : a journal of neurology, 137, 12-32. doi:10.1093/brain/awt162

Meister, M. L. R., Hennig, J. A., & Huk, A. C. (2013, Feb). Signal multiplexing and single-neuron computations in lateral intraparietal area during decision-making. J Neurosci, 33(6), 2254-2267. doi:10.1523/ JNEUROSCI.2984-12.2013

Michelet, T., Duncan, G. H., & Cisek, P. (2010, July). Response competition in the primary motor cortex: corticospinal excitability reflects response replacement during simple decisions. Journal of neurophysiology, 104, 119-127. doi:10.1152/jn.00819.2009

Mostert, P., Kok, P., & de Lange, F. P. (2015). Dissociating sensory from decision processes in human perceptual decision making. Sci Rep, 5, 18253. doi:10.1038/srep18253

Myers, N. E., Rohenkohl, G., Wyart, V., Woolrich, M. W., Nobre, A. C., & Stokes, M. G. (2015, Dec). Testing sensory evidence against mnemonic templates. Elife, 4. doi:10.7554/eLife.09000

O’Connell, R. G., Dockree, P. M., & Kelly, S. P. (2012, November). A supramodal accumulation-to-bound signal that determines perceptual decisions in humans. Nat Neurosci, 15(12), 1729-1735. doi:10.1038/ nn.3248

O’Regan, J. K., & Noe, A. (2001, October). A sensorimotor account of vision and visual consciousness. The Behavioral and brain sciences, 24, 939-73; discussion 973-1031. doi:10.1017/S0140525X01000115

Park, H., Lueckmann, J.-M., von Kriegstein, K., Bitzer, S., & Kiebel, S. J. (2016). Spatiotemporal dynamics of random stimuli account for trial-to-trial variability in perceptual decision making. Sci Rep, 6, 18832. doi:10.1038/srep18832

Park, I. M., Meister, M. L. R., Huk, A. C., & Pillow, J. W. (2014, Oct). Encoding and decoding in parietal cortex during sensorimotor decision-making. Nat Neurosci, 17(10), 1395-1403. doi:10.1038/nn.3800

Pearson, J. M., Heilbronner, S. R., Barack, D. L., Hayden, B. Y., & Platt, M. L. (2011, April). Posterior cingulate cortex: adapting behavior to a changing world. Trends in cognitive sciences, 15, 143-151. doi:10.1016/ j.tics.2011.02.002

Philiastides, M. G., Heekeren, H. R., & Sajda, P. (2014, Dec). Human scalp potentials reflect a mixture of decision-related signals during perceptual choices. J Neurosci, 34 (50), 16877-16889. doi:10.1523/JNEUROSCI. 3012-14.2014

Philiastides, M. G., Ratcliff, R., & Sajda, P. (2006, August). Neural representation of task difficulty and decision making during perceptual categorization: a timing diagram. The Journal of neuroscience : the official journal of the Society for Neuroscience, 26, 8965-8975. doi:10.1523/ JNEUROSCI.1655-06.2006

Philiastides, M. G., & Sajda, P. (2006, April). Temporal characterization of the neural correlates of perceptual decision making in the human brain. Cereb Cortex, 16(4), 509-518. doi:10.1093/cercor/bhi130

Selen, L. P. J., Shadlen, M. N., & Wolpert, D. M. (2012, Feb). Deliberation in the motor system: reflex gains track evolving evidence leading to a decision. J Neurosci, 32(7), 2276-2286. doi:10.1523/JNEUROSCI.5273-11.2012

Siegel, M., Donner, T. H., Oostenveld, R., Fries, P., & Engel, A. K. (2007, March). High-frequency activity in human visual cortex is modulated by visual motion strength. Cerebral cortex (New York, N.Y. : 1991), 17, 732-741. doi:10.1093/cercor/bhk025

Smulders, F. T. Y., & Miller, J. O. (2012). The lateralized readiness potential. In E. S. Kappenman & S. J. Luck (Eds.), The oxford handbook of event-related potential components. Oxford University Press. doi:10.1093/ oxfordhb/9780195374148.013.0115

Taulu, S., Simola, J., & Kajola, M. (2005, Sept). Applications of the signal space separation method. IEEE Transactions on Signal Processing, 53(9), 3359-3372. doi:10.1109/TSP.2005.853302

Thura, D., & Cisek, P. (2014, Mar). Deliberation and commitment in the premotor and primary motor cortex during dynamic decision making. Neuron, 81 (6), 1401-1416. doi:10.1016/j.neuron.2014.01.031

Wyart, V., de Gardelle, V., Scholl, J., & Summerfield, C. (2012, nov). Rhythmic fluctuations in evidence accumulation during decision making in the human brain. Neuron, 76(4), 847-858. doi:10.1016/j.neuron.2012.09.015

